# Sediment chemistry controls methane emissions from lake littoral zones

**DOI:** 10.1101/2025.07.09.663921

**Authors:** Andrew J Tanentzap, Samuel G Woodman, Olesya Kolmakova, Yi Zhang

**Affiliations:** Ecosystems and Global Change Group, Department of Plant Sciences, University of Cambridge, Cambridge, CB2 3EA, United Kingdom; Ecosystems and Global Change Group, School of the Environment, Trent University, Peterborough, Ontario, K9L 0G2, Canada

**Keywords:** biogeochemistry, lake sediment, methanogenesis, methanotrophy, polyphenolics, redox potential

## Abstract

Methane emissions from the nearshore zones of lakes are relatively large but can vary by several orders of magnitude. Here we compared predictions for how sediment chemistry and microbial communities influenced methane emissions from 19 littoral sites in the UK varying in organic matter sources and microbial composition. Our approach was to compare multiple predictions to explain methane fluxes from sediment chemistry and microbial composition using path analysis. We found that the prediction that organic matter composition, namely the concentration of polyphenolics, controls methane emissions by changing electrochemical conditions to favour certain methanogen taxa was hundreds of times better supported than predictions involving abundances of all methanogens and methanotrophic bacteria, methanogen diversity, or other physicochemical conditions. Diffusive CH_4_ fluxes were estimated to increase by 3.1– to 16.6-times (95% confidence interval) with increasing polyphenolic concentrations, almost entirely because they lower reduction-oxidation potentials that shift methanogen composition towards widespread taxa positively associated with methanogenesis. Rather than strongly inhibiting methane-producing microorganisms, our results suggest polyphenolics change reduction-oxidation potentials to favour acetoclastic and methylotrophic methanogens. These results help explain conflicting evidence about the responses of methane to sediment chemistry and can improve future predictions of aquatic carbon cycling.

**Manuscript Highlights:**

- Polyphenolics predicted nearshore CH_4_ fluxes better than other environmental factors
- CH_4_ fluxes increased with polyphenolics that lowered redox to favour methanogenesis
- We help explain conflicting responses of CH_4_ to variation in sediment chemistry

## Introduction

Freshwaters account for half of all global methane emissions, but there is considerable variation in these emissions among and within freshwaters that limits the accuracy of climate and biogeochemical models (Rosentreter and others 2021). In lakes, most methane production occurs in sediments because the lack of oxygen and the availability of both specific nutrients and electron acceptors, such as nitrate or sulfate, favour the anaerobic respiration of organic matter (OM). Sediment chemistry, such as oxygen and nutrient conditions, should therefore be central to explaining within– and among-site variation in lake methane emissions. However, much of our understanding of how chemical conditions influence methanogenesis comes from peatland soils (Yavitt and others 1997; Hodgkins and others 2014; Fenner and Freeman 2020) and livestock rumens (Min and others 2020), with less known about lake sediments.

Lake sediments should vary widely in chemistry and resulting methane production because they receive different aquatic and terrestrial OM inputs (West and others 2012; Grasset and others 2018). Secondary metabolites, particularly polyphenolics, vary greatly among these inputs and contribute to chemical differences between aquatic and terrestrial OM (Emilson and others 2018). Polyphenolics are electron donors and can reduce reduction-oxidation (redox) potentials that electrochemically favour methanogenesis (Walpen and others 2018; Kane and others 2019). Anaerobic decomposition of complex OM, such as polyphenolics, can also consume electron acceptors and further reduce redox potentials (LaRowe and Van Cappellen 2011), as well as produce methanogenesis precursors, such as acetate (McGivern and others 2021). These processes may explain positive correlations between methane production and both phenols and phenolic precursors (Heslop and others 2017). However, past studies have not tested explicitly the role of redox potential and its interaction with polyphenolics in explaining variation in methane emissions.

Polyphenolic concentrations associated with different OM sources also influence methane production by altering microbial composition and function (Yakimovich and others 2020). Some studies have found polyphenolic concentrations correlated negatively with methane concentrations because they were toxic to methanogens and/or inactivated enzymes involved in OM decomposition (Emilson and others 2018), such as by binding nutrients required for oxidation of phenolics (Fenner and Freeman 2020; Ding and others 2022). One explanation for the discrepancy among studies in the response of methane production to polyphenolics is that redox potentials may have been low enough in the study reporting a positive correlation that few electron acceptors competed with acetoclastic methanogenesis (Heslop and others 2017). At high redox potentials, acetoclastic methanogens, such as in the genera *Methanosarcina* and *Methanosaeta*, may decline in abundance because they are outcompeted for shared substrates, such as H_2_, by other microbes, e.g. sulfate-reducing bacteria (Conrad and others 1987; Zhuang and others 2018). Therefore, redox potential likely mediates the effect of polyphenolics on methane production by influencing methanogen community composition and the electrochemical favourability of methanogenesis.

Many other factors besides sediment chemistry influence methane fluxes from lake sediments but their importance relative to sediment chemistry remains poorly understood. These factors include the presence and composition of aquatic vegetation, which can directly transfer methane to the atmosphere through porous tissue known as aerenchyma (Bodmer and others 2024; Desrosiers and others 2022), in addition to producing OM with relatively low polyphenolic concentrations (Emilson and others 2018). Microbial community composition and activity, including of both methanogens and methanotrophs that are central to determining methane fluxes, vary among lakes with different physicochemical conditions (Bertolet and others 2019; Yakimovich and others 2020; Winder and others 2023). For example, acidic pH can increase the thermodynamic favourability of hydrogenotrophic methanogenesis but not acetoclastic methanogenesis (Jin and Kirk, 2018), whereas a wide range of pH can enable oxidation by methanotrophs (Chowdhury and Dick, 2013). The activity of these communities also depends highly on temperature (Yvon-Durocher and others 2014; DelSontro and others 2016), which itself varies across environmental gradients (Deemer and Holgerson 2021). The impact of warmer temperatures on methane fluxes also varies with season, depth, and landscape (Marotta and others 2014; West and others 2016; Deemer and Holgerson 2021), and all these factors can influence the contribution of ebullition (McGinnis and others 2006; DelSontro and others 2016). For example, warmer temperatures can decrease the solubility of methane in pore water, thereby releasing more bubbles from sediment to the lake surface (Aben and others 2017). Finally, shallower water columns can allow methane to escape oxidation as it travels from sediments to the lake surface (West and others 2016) but also allow more oxygen to penetrate sediments and increase aerobic methane oxidation (Li and others 2021).

Here we determined the importance of sediment chemistry versus other environmental factors in explaining variation in methane emissions from lake littoral zones. We focused on littoral zones because methane fluxes are highest in their warm and shallow waters and can vary by several orders of magnitude even therein (Juutinen and others 2003; Bastviken and others 2008; Desrosiers and others 2022). We related methane fluxes from 19 UK sites to sediment chemistry, abundances of methane-producing and methane-oxidising microbes, and methanogen composition. To test the overarching hypothesis that sediment chemistry, specifically polyphenolic concentrations and redox potential, explains variation in methane emissions, we first compared macrophyte beds to littoral deposition zones. Sediments in these sites should differ in chemical composition, especially polyphenolic concentrations (Emilson and others 2018), because they receive primarily aquatic versus terrestrial OM, respectively. We then used path analysis to test multiple explicit causal mechanisms for how sediment chemistry influences methane emissions. This approach allowed us to disentangle the direct effects of sediment chemistry on methanogenesis, such as through polyphenolics changing redox potential or inactivating enzyme activity, from indirect effects, such as shifting microbial community composition. Together, our approach offers new insights into how methane emissions from lake littoral zones concurrently vary with physicochemical conditions and microbial communities.

## Methods

### Study sites

We sampled 19 sites across northern England and Scotland from 3-8 September 2018 (Table 1). Sampling coincided with late-summer when temperatures, and thus methanogenesis, peaks (Delwiche and others 2021). All sites were littoral with ca. 1 m of overlying water to control effects of depth (West and others 2016), and either included or excluded emergent macrophytes (n = 6 and 13, respectively). Sites without macrophytes were nearshore deposition zones associated with channelized inflows to capture variation in OM inputs. Catchments for these inflows were dominated by acid and heather grasslands (Table 1; Supplementary Methods S1).

**Table 1.** Characteristics of littoral study sites in the northern UK. Latitude (Lat) and longitude (Lon) correspond to the sampling location of each of 19 littoral sites. We compared sites with macrophyte vegetation versus the littoral deposition zones of lake inflows, which we expected to differ in their OM inputs. Two distinct deposition zones from separate inflows were sampled in two lakes (Lochy and Stack) rather than sampling additional lakes. At each site, we estimated mean (± standard error) for diffusive and ebullitive CH_4_ fluxes from sediment, measured polyphenolic concentrations in sediment expressed as gallic acid equivalents (GAE) per g dry weight of sediment, and measured redox potential standardized to pH 7 and specific ultraviolet absorbance at 254 nm (SUVA) in sediment porewater.

| Lat | Lon | Lake | Lake area (km<sup>2</sup>) | Mean lake depth (m) | Diffusive CH<sub>4</sub> (μmol C m<sup>-2</sup> hr<sup>-1</sup>) | Ebullitive CH<sub>4</sub> (μmol C m<sup>-2</sup> hr<sup>-1</sup>) | Total CO<sub>2</sub> (mmol C m<sup>-2</sup> hr<sup>-1</sup>) | Polyphenolics (mg GAE g<sup>-1</sup> sediment) | Redox potential (mV) | SUVA (L mg C<sup>-1</sup> m<sup>-1</sup>) |
| --- | --- | --- | --- | --- | --- | --- | --- | --- | --- | --- |
| *Sites with macrophyte vegetation* |  |  |  |  |  |  |  |  |  |  |
| 56.667 | -5.033 | Achtriochtan | 0.1 | NA | 504.3 (3.3) | 0.00 (0.00) | 3.1 (0.1) | 0.14 | 165.3 | 0.18 |
| 58.095 | -4.974 | Awe | 0.3 | 3.9 | 384.5 (0.8) | 0.00 (0.00) | 5.2 (0.1) | 1.48 | -102.5 | 3.17 |
| 56.608 | -4.755 | Ba | 1.9 | 1.9 | 395.5 (1.0) | 0.00 (0.00) | 2.3 (<0.1) | 0.25 | -32.9 | 5.04 |
| 58.370 | -5.078 | Claise | 0.1 | NA | 409.7 (2.8) | 181.9 (2.4)<sup>a</sup> | 7.5 (0.2) | 1.75 | -96.5 | 3.33 |
| 56.671 | -4.001 | Kinardochy | 0.2 | 4.3 | 439.5 (1.3) | 0.00 (0.00) | 0.5 (0.1) | 1.47 | -111.7 | 3.00 |
| 58.323 | -4.911 | Stack | 2.2 | 9.9 | 267.7 (1.1) | 0.00 (0.00) | 3.3 (<0.1) | 0.71 | -97.4 | 5.70 |
| *Sites without macrophyte vegetation* |  |  |  |  |  |  |  |  |  |  |
| 58.304 | -4.902 | Allt Ceann | 3.4 | 17.4 | 141.5 (3.3) | 137.6 (6.4)<sup>b</sup> | -23.1 (0.2) | 0.17 | -77.1 |  |
| 58.151 | -4.982 | Assynt | 7.1 | 14.6 | 286.0 (0.8) | 0.00 (0.00) | 3.1 (0.1) | 0.02 | 193.5 | 5.34 |
| 54.669 | -3.241 | Bassenthwaite | 5.0 | 12.6 | 656.3 (2.5) | 0.00 (0.00) | 20.9 (0.1) | 0.41 | -55.0 | 2.36 |
| 56.992 | -4.864 | Lochy | 16.2 | 3.2 | 139.3 (0.7) | 0.00 (0.00) | 1.3 (0.1) | 0.07 | 34.6 | 3.69 |
| 57.025 | -4.825 | Lochy | 16.2 | 3.2 | 1531.3 (40.9) | 694.2 (118.7)<sup>b</sup> | 104.3 (1.9) | 0.69 | -136.2 | 0.20 |
| 56.147 | -4.659 | Lomond | 68.5 | 34.0 | 86.1 (2.7) | 97.1 (10.9)<sup>a</sup> | 0.8 (0.4) | 0.92 | -36.3 | 2.41 |
| 56.293 | -4.294 | Lubnaig | 2.0 | 15.4 | 151.4 (4.4) | 0.00 (0.00) | 3.2 (0.8) | 0.09 | 167.4 | 6.89 |
| 57.331 | -4.452 | Ness | 51.6 | 61.9 | 1491.8 (2.2) | 0.00 (0.00) | 26.6 (0.1) | 0.13 | -59.8 | 4.97 |
| 58.145 | -4.605 | Shin | 31.5 | 10.1 | 92.8 (0.5) | 0.00 (0.00) | 1.3 (0.1) | 1.36 | -67.2 | 1.75 |
| 58.345 | -4.957 | Stack | 2.2 | 9.9 | 38.6 (1.1) | 71.6 (6.2)<sup>b</sup> | 18.7 (14.3) | 0.21 | 11.5 | 5.28 |
| 58.334 | -4.929 | Stack | 2.2 | 9.9 | 16.1 (0.6) | 0.00 (0.00) | 4.0 (1.5) | 0.05 | 142.2 | 2.79 |
| 56.526 | -4.120 | Tay | 24.1 | 31.0 | 21.7 (0.6) | 0.00 (0.00) | 1.2 (0.2) | 0.06 | 212.4 | 0.98 |
| 54.569 | -2.930 | Ullswater | 8.0 | 18.2 | 25.1 (5.2) | 32.1 (2.7)<sup>a</sup> | -9.4 (2.7) | 0.02 | 139.7 | 2.48 |

<sup>a</sup>one ebullition event; <sup>b</sup> two distinct ebullition events

### Greenhouse gas fluxes

Daytime CH_4_ and CO_2_ fluxes at the air-water interface were measured using a floating plastic chamber (volume = 27.6 L, area = 0.08 m^2^) covered in reflective tape to minimise heating. The chamber was deployed from nearshore with a 5 m telescopic pole and we ensured that it did not cover any emergent macrophyte vegetation. The chamber was connected in a closed loop to an off-axis integrated cavity output spectrometer (ultraportable greenhouse gas analyzer 915, Los Gatos Research, USA) that recorded greenhouse gas (GHG) concentrations every second for 10 mins once per site (Gonzalez-Valencia and others 2014; Kuhn and others 2021). Because of battery failures, two sites could not be measured for the entire 10 mins and had shorter readings (6 and 8.5 mins). We acknowledge GHG fluxes vary diurnally and among days (Sieczko and others 2020), but our aim here was to test environmental predictors concurrent with fluxes rather than reconstruct seasonal or annual carbon budgets. Any associations between GHG fluxes and their environmental predictors are likely to be consistent at different times even if the absolute values differed. The measured fluxes were then normalised to hourly rates so that they could be directly compared between lakes and with environmental predictors. Although our measurement periods were relatively short, they were identical in duration to other published studies (Grasset and others 2016; Barbosa and others 2021; Peacock and others 2021), and were arguably more direct measurements of fluxes than those derived by applying gas exchange velocities to dissolved gas concentrations (Holgerson and Raymond 2016). Importantly, because rates of diffusive fluxes were estimated from the slope of linear changes in methane concentrations over time (see paragraph below), they should largely be invariant with the duration over which they were measured when temperature, wind, and rain conditions remain stable (Natchimuthu and others 2016; Sø and others 2024). As weather during each measurement period was consistent, rates of diffusive fluxes should primarily reflect sediment characteristics at the time of measurement.

To generate total flux estimates, we multiplied rates of diffusion and ebullition by their estimated duration, summed the values, and divided by the length of the entire measurement period. Diffusive fluxes were calculated for each site from the slope of a linear regression between GHG concentrations and measurement time, and corrected for air temperature and atmospheric pressure using the ideal gas law (Erkkilä and others 2018; Peacock and others 2021). As rates of diffusive fluxes will be biased upwards by fitting a regression to observed concentrations in the presence of ebullition events, because ebullition causes sudden, non-linear increases in GHG concentrations, we used an established statistical approach that identifies ebullition events nearly identically to sudden changes in gas transfer velocities (Goodrich and others 2011; Barbosa and others 2021). We first visually inspected each linear regression to identify any sudden, non-linear spikes in concentration lasting <2 min, after which concentrations continued to climb at rates similar to before the spikes occurred. We then estimated separate slopes (i.e. fluxes) for these putative ebullition events by fitting piecewise linear regressions with estimated breakpoints using the segmented v.1.6-4 package (Muggeo 2017) in R v4.0 (R Core Team, 2022). We used the slope estimates at the estimated breakpoints to calculate ebullition rates after Goodrich and others (2011). All the regression models were statistically significant and fitted the data well for CH_4_ and CO_2_; median (inter-quartile range) for R^2^ = 0.990 (0.990 to 0.999) and 0.946 (0.865 to 0.975), respectively. Finally, we used the ebullition rates to separate diffusive fluxes for our statistical analyses rather than solely use total fluxes, which will potentially be biased by the ability to capture ebullition in some sites but not others during our relatively short measurement period.

### Physicochemical conditions

We sampled porewater and sediment directly beneath areas used for flux measurements immediately after recording GHGs. Sediment porewater was collected with non-destructive micro-tensiometers that representatively capture dissolved components at the sediment-water interface (Seeberg-Elverfeldt and others 2005). We inserted two 12 cm long MacroRhizons (0.15 µm pore size; Rhizosphere Research Products, the Netherlands) per site at a 45° angle into sediment. The end of the MacroRhizons was connected via tubing to a 60 mL syringe above the water surface. Tubing and syringes were purged twice before collecting porewater. 20 mL of sediment porewater was used immediately to measure water temperature, pH, and redox with a handheld multiprobe (HI-991002, Hanna Instruments, USA) paired with a gel-filled electrode (HI-1297) and platinum redox sensor silver/silver chloride internal reference system that were calibrated before measurements. The probe reports redox in mV with an accuracy of ± 2 mV and values were standardized to pH 7 by subtracting or adding 58 mV for every measured pH unit (accuracy: ± 0.02 pH) beneath or above 7, respectively (Wetzel 2001). We then passed water through pre-rinsed 0.2 µm cellulose acetate filters (Sartorius AG, Germany) into 20 mL glass vials with no headspace that were kept in the dark at 4°C for up to a week for dissolved organic carbon (DOC) and OM quality analyses. Finally, we collected surface sediment to a depth of 7.5 cm using a modified piston corer (2.6 cm diameter). Sediment was extruded into 50 mL sterile centrifuge tubes and immediately flash frozen in liquid nitrogen. Sediment was homogenized before microbial DNA extraction and sediment chemistry analyses by mixing samples with a sterile spatula. We did not collect multiple samples from a site (i.e. within-site variation) as our goal was not to reconstruct chemistry across the entire littoral zone. Rather, we wanted to capture differences in physiochemical conditions and microbial communities during flux measurements.

In the lab, we measured porewater and sediment chemistry. Porewater DOC concentrations were measured using wet chemical breakdown coupled with spectrophotometry after purging samples of existing dissolved inorganic carbon (LCK385 kit, Hach Lange, Germany). To estimate OM quality, we measured absorbance at 254 nm divided by DOC concentrations to calculate specific ultraviolet absorbance (SUVA). Larger SUVA values indicate more aromatic, high molecular weight compounds (Lavonen and others 2015). Sediments were air-dried and polyphenolics were extracted in triplicate per site by shaking 0.15 g of dried material in 1.5 mL of methanol acidified with 1.5 mL of 0.12M methanesulfonic acid for 1 hour at 23°C followed by incubation at 20°C for 22 hours (Halvorson and others 2009). Polyphenolic concentrations were determined with colorimetry using the Folin-Ciocalteau method and averaged across replicates per site in gallic acid equivalents (Ainsworth and Gillespie 2007). We also measured OM content as weight loss on ignition for 7 hours in a 500°C muffle furnace.

### Microbial functional gene abundance

We used qPCR to quantify copy numbers of methanogen and both aerobic and anaerobic methanotroph functional genes in sediment. DNA was extracted from 0.25 g of freeze-dried sediment using a Powersoil DNA Isolation Kit (MO BIO Laboratories, USA) following the manufacturer’s protocol. We amplified the methyl coenzyme M reductase (*mcrA*) gene to quantify the abundance of methanogens, and methylocella (*mmoX*) and particulate methane monooxygenase (*pmoA*) genes to quantify abundances of methane-oxidizing bacteria. For *mcrA*, we targeted the mlas-F and mcrA-rev primer pair (Angel and others 2012). We targeted the mmoXLF and mmoXLR primers for *mmoX* (Rahman and others 2011), and different primer pairs for type Ia (A189f/Mb601r), type Ib (A189f/Mc468r), and type II (A189f/A621r) versions of *pmoA* genes (Putkinen and others 2018). The qPCR was conducted in triplicate for each sample with a HOT FIREPol® EvaGreen® qPCR Supermix (Soils BioDyne, Estonia) on a CFX96 Touch real-time PCR detection system (Bio-Rad Laboratories, USA) with minor modifications (Supplementary Methods S1). Triplicate Ct values were averaged and converted to copy number using the standard curve of each target gene and expressed per gram dry weight of sediment. A standard curve for absolute copy number of each gene (R^2^ = 0.980-0.996) was generated by serial dilution of bands purified after PCR amplification (efficiency = 73-111%) of sediment DNA extracts with a Monarch® DNA Gel Extraction Kit (New England Biolabs, USA). Purified DNA concentrations were measured with a Qubit™ dsDNA high sensitivity assay kit (Life Technologies, UK), and copy numbers determined given target gene length.

### Methanogen community composition

We characterised methanogen composition in each sample by amplicon sequencing the *mcrA* gene. We added 1 to 10 ng of extracted DNA to PCR reactions with 1X LongAmp Taq2X MasterMix (New England Biolabs, USA) and 0.4 mM primers mlasF (5′-GGT GGT GTM GGD TTC ACM CAR TA-3′) and mcrA-rev (5′-CGT TCA TBG CGT AGT TVG GRT AGT-3′) tailed with standard Oxford Nanopore Technologies (ONT) sequences. The thermocycling conditions were: initial denaturation for 1 min at 94°C; 5 cycles of 30 s at 94°C, 30 s at 60°C, 50 s at 65°C; 30 cycles of 30 s at 94°C, 30 s at 62°C, 50 s at 65°C, and final elongation at 65°C for 5 min. The amplification product was purified by solid-phase reversible immobilization on purification beads (Illumina, USA) and used as a template for barcoding amplification with the PCR Barcoding Expansion Kit (EXP-PBC001, Oxford Nanopore Technologies, UK). DNA repair, end-prep, adapter ligation, and clean-up were performed using the Ligation Sequencing Kit (SQK-LSK110, Oxford Nanopore Technologies, UK) and NEBNext® Companion Module for Oxford Nanopore Technologies® Ligation Sequencing (E7180S, New England Biolabs, USA) according to the manufacturer’s protocol.

The 19 samples were sequenced on ONT Flongle flow cells (FLO-FLG001, R9.4.1 chemistry) with a MinION to generate ca. 1 million reads. Raw reads were basecalled and demultiplexed with Guppy v6.3.8. FastQC v.0.11.9 was used to assess read quality, and reads with a Phred score <10 were removed during basecalling, which is relatively conservative for ONT data (Rodríguez-Pérez and others 2021; Sereika and others 2022). We further filtered reads to between 400 and 600 bp to capture high-quality fragments that were close in read length to the expected fragment size while accounting for variation in read length due to sequencing variability or adapter trimming. Reads were then mapped to a taxonomic database of *mcrA* sequences that supports species-level identification (Yang and others 2014) using Minimap2 (Li 2018), retaining those with a mapping quality >0 and alignment score ≥300. Overall, we retained between 17,318 to 92,371 reads per sample and all samples in our subsequent analyses (Table S1).

We estimated methanogen diversity at the species level using the Shannon index that accounts for the relative abundances of taxa, estimated by normalising the read counts in each sample to sum to one. We avoided adjusting sequencing depth prior to comparing diversity as this approach can unnecessarily discard data (McMurdie and Holmes, 2014; but see Schloss, 2024). Importantly, this decision did not change any of our conclusions. We calculated the mean Shannon index after randomly resampling communities 100 times in each sample to the lowest number of species recovered in a single sample across the dataset. Sample-specific diversity estimated with and without rarefaction was highly correlated (Pearson correlation r = 0.99997, p <0.001), as was the correlation between the Shannon index and the log-transformed inverse Simpson index (r = 0.98, p <0.001), which can be less sensitive to sampling effort (Schloss 2024).

We separately identified archaeal anaerobic methanotrophs (ANMEs) by mapping reads with the same procedure as for *mcrA*. We used a custom database compiled from all matches (n = 207) in the NCBI Nucleotide database to the search terms “ANME AND mcrA” and the accessions in Kevorkian and others (2021).

### Statistical analyses comparing macrophyte beds and littoral deposition zones

We tested if polyphenolic concentrations and CH_4_ fluxes differed, on average, between macrophyte beds and littoral deposition zones using *t*-tests. We used the Welch approximation to calculate degrees of freedom where variances between groups were unequal according to Levene’s test for homogeneity of variance. Non-normally distributed, positive variables were log-transformed in all analyses. Rather than report p-values, we calculated 95% confidence intervals (CIs) of effect sizes, i.e. to summarise how much larger values were in one treatment versus another. Because redox potential was non-normally distributed and could not be log-transformed as it included negative values, we compared it between sites with an exact Wilcoxon-Mann-Whitney test. Analyses were undertaken in R v4.0 (R Core Team, 2022).

To compare microbial community composition between macrophyte beds and littoral deposition zones, we used permutational analysis of variance (PERMANOVA) with the adonis2 function in the R package vegan v.2.6 (Oksanen and others 2022). We estimated compositional dissimilarity with the Bray-Curtis index with data aggregated at the Family-level and calculated the effect of site type with 999 random permutations. We also tested the amount of variance in diffusive CH_4_ fluxes explained by the entire community composition matrix using partial least squares regression (PLSR) fitted with the R package pls v.2.8 (Mevik and Wehrens 2007). We restricted the predictors to the 96 taxa that reached >1% relative abundance in at least one sample. We then calculated variance in projection (VIP) scores for each explanatory variable (i.e. taxa), with VIP scores ≥ 1 considered to be highly predictive of the CH_4_ fluxes (Chong and Jun 2005). We also tested how the relative abundance of individual methanogens varied with CH_4_ fluxes using Spearman rank correlations. We restricted this analysis to methanogens present in most (i.e. >50%) sites.

### Path analysis to predict CH_4_ fluxes

We compared 18 explanations for how diffusive CH_4_ fluxes varied with sediment chemistry, environmental conditions, and microbial communities (Table S2) using path analysis fitted with the piecewiseSEM v.2.3 package in R (Lefcheck 2016). Path analysis involves specifying hypothesised cause-and-effect linkages estimated by interconnected regression models.

Our overarching hypothesis that sediment chemistry explains CH_4_ fluxes leads to the prediction (*P*) that higher polyphenolic concentrations arise from increased sediment OM, which reflect terrestrial inputs (Emilson and others 2018), and directly influence methanogenesis by lowering redox potential. Lower redox potential can directly change the (*P1*) electrochemical favourability of reactions involved in methanogenesis (Kane and others 2019; Walpen and others 2018) and (*P2*) shift methanogen composition in ways that influence their activity. To test (*P2*), we summed the relative abundance of individual methanogen sequences present in a majority of samples (i.e. found in >50% of sites) and that had a positive Spearman rank correlation (*p* < 0.05) with diffusive CH_4_ fluxes. This aggregation allowed us to test explicitly if the effect of redox potential on CH_4_ was mediated by the composition of taxa most associated with methanogenesis. Because the efficiency of methanogenesis varies among taxa (e.g. Xie and others 2024), these compositional effects can be a better predictor of CH₄ emissions than the total abundance of all methanogens. Although we could also model how community composition influenced CH_4_ using an approach like PLSR (e.g. Emerson and others 2021), the path analysis could not then model the abundances of thousands of individual taxa contingent on other variables, such as polyphenolic concentrations. Instead, we also explored replacing the aggregated variable with abundances of individual taxa with VIP scores ≥ 1 in our separate PLSR. Finally, we tested if higher polyphenolic concentrations (*P3*) supply substrates for methanogenesis and/or inactivate associated enzymes by including a direct path in our model from polyphenolic concentrations to CH₄ emissions (Heslop and others 2017; Fenner and Freeman 2020).

Sediment chemistry can also indirectly influence CH_4_ fluxes by changing microbial abundance and composition. Thus, we also tested if methanogen abundances decreased with polyphenolic concentrations (Emilson and others 2018), alongside the influence of known chemical controls: pH, DOC concentration, and SUVA of sediment porewater (Lew and Glińska-Lewczuk 2018; Bertolet and others 2019; Yakimovich and others 2020). Total methanogen abundances can influence CH_4_ (*P4*) directly, such if greater numbers increase total activity, and (*P5*) indirectly by influencing methanogen diversity, because larger communities are more likely to include more species simply by chance. A greater diversity can subsequently enhance CH_4_ fluxes because different methanogen species can utilise complementary substrate and metabolic pathways, leading to higher overall CH_4_ production. Methanogen diversity was also modelled from pH, DOC, and SUVA. Finally, CH_4_ fluxes can (*P6*) depend on abundances of methane oxidisers, resulting in covariation between CH_4_ and CO_2_ fluxes. We separately modelled abundances of methanotrophs that used *mmoX* or *pmoA* with the same predictors we used for methanogens.

Each prediction (*P1*-*P6*) was varied in two additional ways. First, we allowed (*P7*-*P12*) both OM and sediment polyphenolic concentrations to co-vary simply because they were associated with separate, unmeasured processes. Second, we allowed (*P13*-*P18*) all of OM, polyphenolics, and redox to co-vary for the same reasons. Throughout, we accounted for variation in CH_4_ and CO_2_ fluxes independent of sediment chemistry and because of water temperature during sampling and the regional climate, such as differences in long-term temperature, solar radiation, and seasonality, and all of which can be approximated by latitude. We used Fisher’s C to test if the overall structure of each model fitted the data and used directed separation tests to identify missing linkages between variables (Grace and Irvine 2020). These latter tests necessitated adding a linkage from OM quality (i.e. SUVA) to polyphenolic concentrations (*P1*-*P6* only), from sediment OM concentration to methanogen and methanotroph abundance, and covariation between methanogen and methanotroph abundances.

Because of high multi-collinearity, we tested *P1*-*P18* separately and compared support for each prediction with the Akaike information criterion corrected for small sample sizes (AICc). This approach of comparing explicit *a priori* predictions is preferable to both model dredging from a single “full” model with all potential predictors and model averaging (see Grace and Irvine 2020 for further discussion). For the best supported models, i.e. lowest AICc, we calculated 95% CIs for standardised effects adjusted for multi-collinearity from 1,000 bootstrapped estimates with the semEff v.0.6.1 R package (Murphy 2023).

## Results

### Patterns of methane fluxes and microbial community composition

Total CH_4_ fluxes ranged from 3.9 to 534.1 µmol C m^-2^ hr^-1^, covering the range observed globally in lakes (Rosentreter and others 2021), and varied with emergent macrophyte vegetation that could export unique organic matter to sediment. Most of the total CH_4_ fluxes were diffusive emissions, with ebullition detected at only six sites potentially because of the relatively short measurement duration (Table 1). We found total CH_4_ fluxes were 1.0-to 5.9-times higher (95% CI estimated from t-tests) in sites with versus without emergent macrophyte vegetation, with diffusive fluxes 1.2-to 7.5-times larger under macrophytes (Table 1). These differences corresponded with 1.3-to 16.4-times higher (t-test 95% CI) sediment polyphenolic concentrations in sites with versus without macrophytes (Table 1). Although redox potential was similar between sites with versus without macrophytes (95% CI for difference in the mean value between sites from t-test: –242 to 24 mV), it strongly negatively correlated with sediment polyphenolic concentrations (Spearman correlation ρ = –0.82, *p* <0.001). Polyphenolic concentrations ranged from 0.02 to 1.75 mg gallic acid equivalent g^-1^ sediment across our sites corresponding with redox potentials of –136 to 212 mV (Table 1).

We found some evidence that methane emissions co-varied with methanogen community composition. We mapped 723,061 mcrA reads retained after quality control to 1,795 unique amplicon sequence variants across our study sites. The unique sequences that we could identify in our samples included at least 5 known orders, 8 families, and 11 genera (Fig. 2, Fig. S1). Of the 52 sequences that could be mapped to at least Family-level, composition was dominated by Methanosarcinaceae, which primarily use acetoclastic and methylotrophic pathways, with far fewer hydrogenotrophic taxa (Fig. 2a, b). Sites with macrophytes had more acetoclastic Methanosaetaceae (Fig. 2a, b), thus differing in community composition (PERMANOVA, F_1,18_ = 9.5, p = 0.005, R^2^ = 0.36). Sixty mapped sequences were found in all samples but none of these could be identified beyond Archaea, highlighting the potentially large number of ubiquitous and novel taxa in lake sediments. Two of these ubiquitous taxa (Genbank accession numbers: FR725517 and AM746839) could be found at high (i.e. > 10%) relative abundance in some sites. Overall, methanogen community composition explained about 67% of the variance in diffusive CH_4_ fluxes based on a PLSR model, with VIP scores between 1.0 and 7.8 for six taxa identifiable only to Archaea (Genbank accession numbers in order of ascending VIP: GU085068, FR725772, AF525519, AB570064, GQ390077, FR725517). Of taxa that occurred in at least 50% of the samples, 57 and 17 correlated positively and negatively with diffusive CH_4_ fluxes, respectively, but none could be identified beneath Order-level (Fig. 2c).

**Figure 1.**
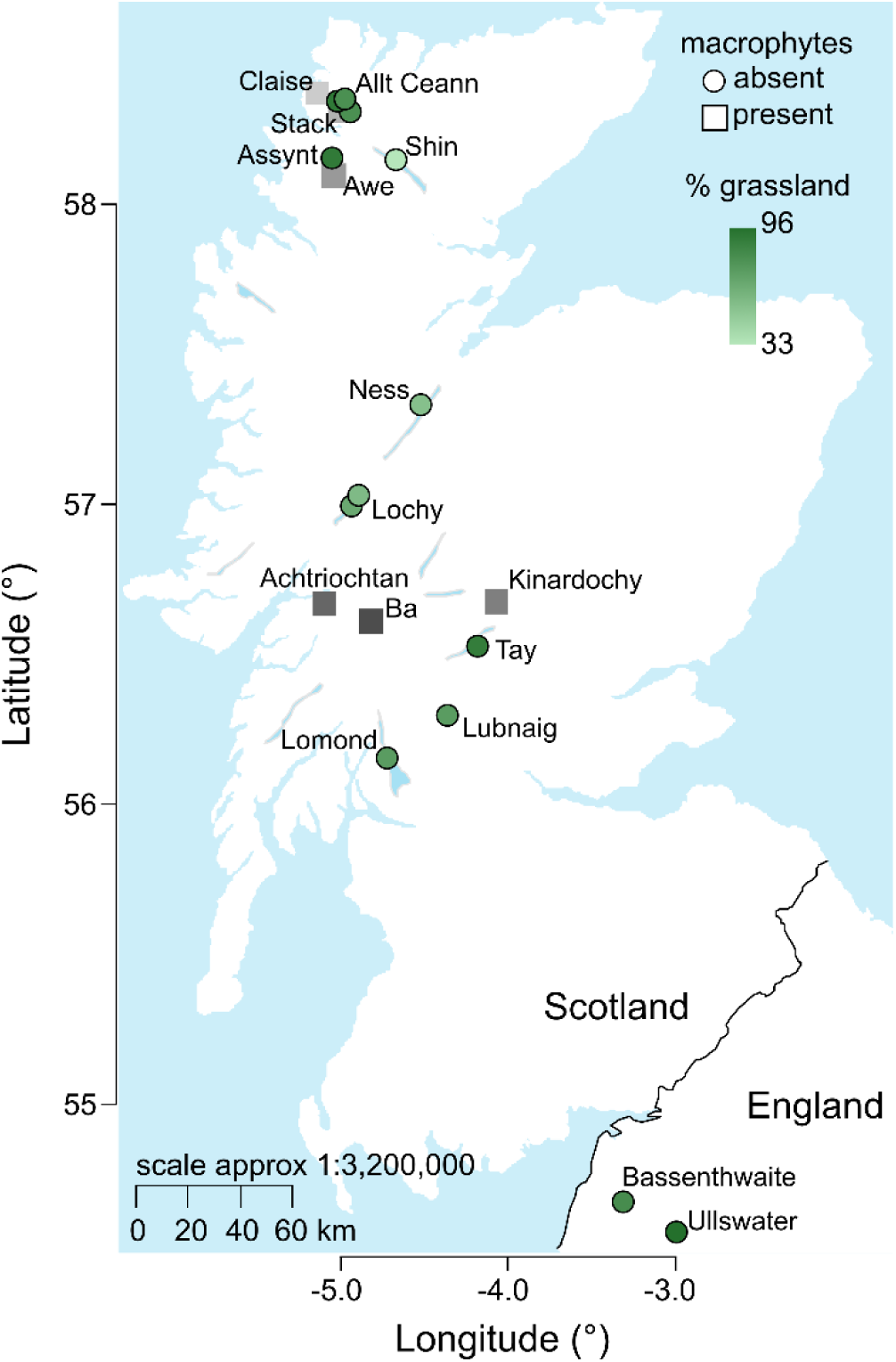
Littoral sediments sampled across northern UK lakes varied in the composition of organic matter inputs. Sites either had emergent macrophyte vegetation present (n=6) or absent (n=13). Sites without macrophyte vegetation were located at the deposition zone of lake inflows that drained catchments with varying grassland vegetation, with darker green indicating a greater percentage of grassland land cover. Grassland vegetation was not plotted for macrophyte sites as these were not located in catchment deposition zones. For sites with macrophytes, we coloured points according to latitude. The figure was generated in R 4.2 using the maps 3.4.1 package with data freely available for reuse from naturalearthdata.com.

**Figure 2.**
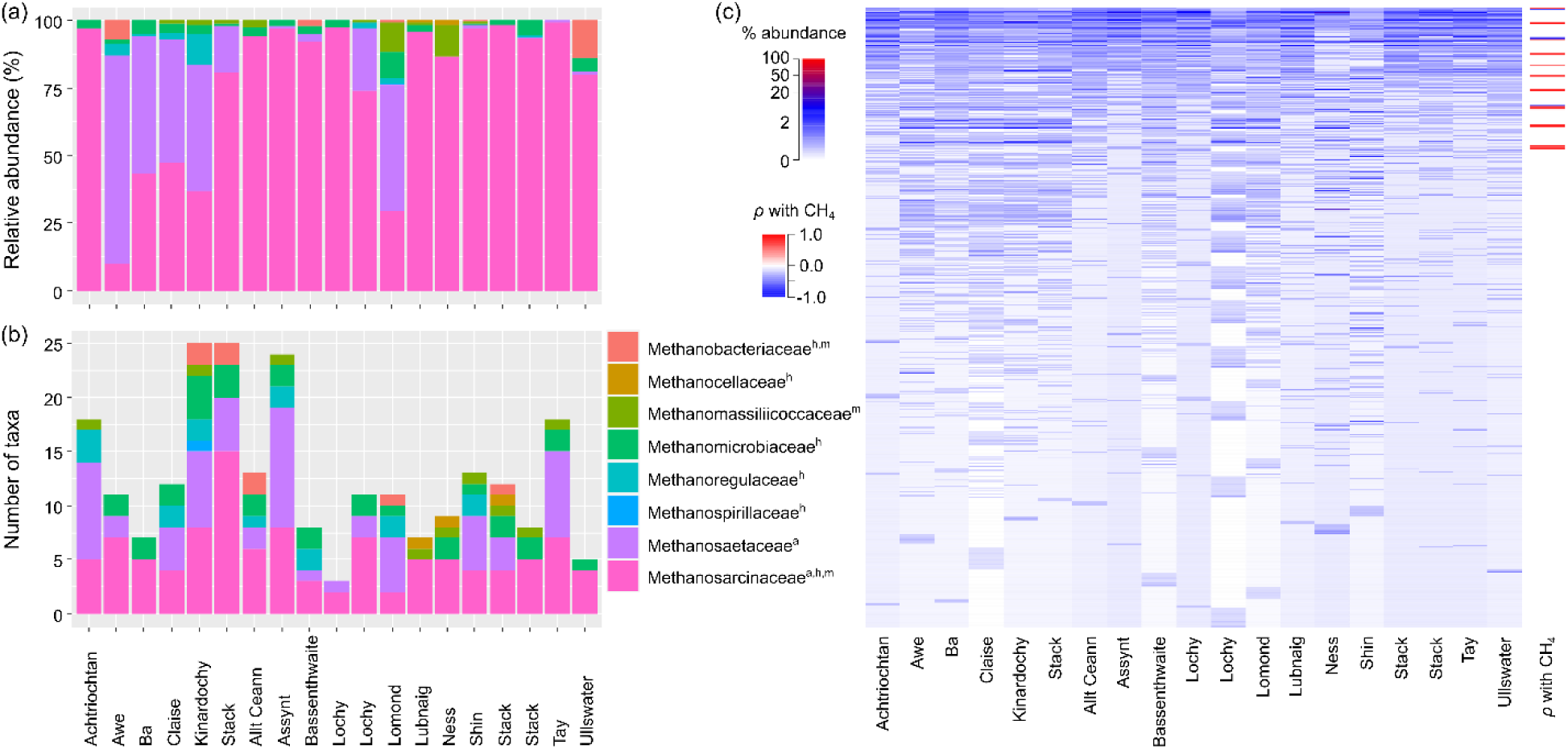
Methanogen composition in littoral lake sediments. (a) Relative abundance of reads and (b) number of unique taxa that could be mapped to the Family-level in each of 19 sites across the northern UK. Families were classified into whether they can undertake ^a^acetoclastic, ^h^hydrogenotrophic, or ^m^methylotrophic methanogenesis (after Liu and Whitman 2008; Ellenbogen and others 2023). (c) Relative abundance across all reads (n = 1,795) mapped to a *mcrA* sequence in each site. Rows are individual taxa sorted from those present in the most (top) to fewest (bottom) sites, with colours corresponding to relative abundance. We also correlated the relative abundance of each taxon with diffusive CH_4_ fluxes across sites and plotted correlations that were statistically significant (p < 0.05). NCBI Genbank accession numbers corresponding to each row are given in Data S1.

We found little evidence of ANMEs (Table S3). Only 1 taxon from a study of ANMEs was present in all samples, but it could only be identified taxonomically as archaea rather than an ANME specifically. Three other taxa, corresponding to a *Candidatus* Methanoperedens sp. and two other potential ANME-2d species, were found in 2 sites with relatively high methane emissions (Bassenthwaite and Ness; Table 1), but with negligible reads (Table S3).

### Sediment chemistry predicts littoral methane fluxes

We found evidence that suggested sediment chemistry influenced CH_4_ emissions through electrochemical conditions rather than by inhibiting methanogens. The path analysis model *P2* that predicted diffusive CH_4_ fluxes indirectly from polyphenolic concentrations had overwhelming support in the candidate set (AIC weight = 98.5%, Table S2; Fig. 3). This model estimated that diffusive CH_4_ fluxes strongly increased with higher polyphenolic concentrations (Fig. 4a), because higher polyphenolic concentrations reduced redox potential (Fig. 4b), which shifted methanogen composition (Fig. 4c). Specifically, lower redox potential shifted methanogen composition towards individual taxa that were widespread, i.e. present in >50% of samples, and positively correlated with diffusive CH_4_ fluxes (Fig. 2c). The summed abundance of these taxa was subsequently tightly coupled to CH_4_ emissions (Fig. 4d). Replacing the summed abundance of these taxa with the individual abundances of the six taxa with VIP scores ≥1 identified by the PLSR analysis did not result in a better supported model as AICc increased by 47.1. A next best-supported model (*P1*) predicted a direct negative effect of redox potential on CH_4_ fluxes, with redox decreasing with increasing polyphenolic concentrations (AIC weight = 0.8%). There was also little evidence that increasing polyphenolics directly promoted CH_4_ fluxes (AIC weight = 0.2%), such as if they provided substrates for methanogenesis (*P3*). Thus, the polyphenolics created highly reduced, anaerobic conditions that favoured certain methanogens (Fig. 3). Diffusive CH_4_ fluxes predicted across the models *P1* to *P3* and weighted by their AIC support were consequently estimated to increase by a mean (95% CI) of 6.2-times (3.1-to 16.6-times) over the observed range of polyphenolic concentrations in sediments (Fig. 4a).

**Figure 3.**
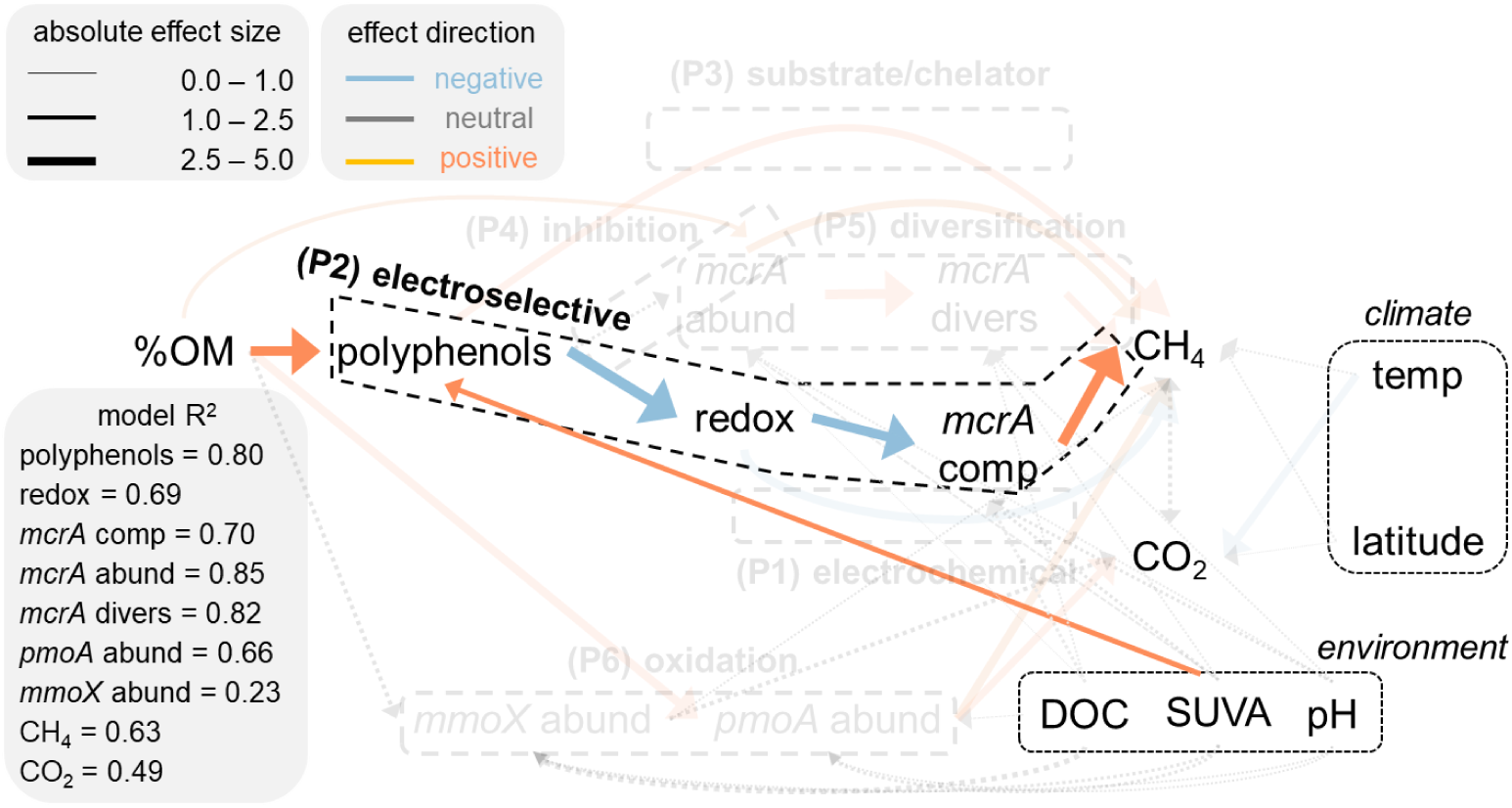
Sediment chemistry mainly controls littoral greenhouse gas emissions. We tested six predictions (*P1*) to (*P6*) shown by broken boxes to explain diffusive CH_4_ and CO_2_ fluxes in 19 littoral sites. Predictions were separately modelled using path analysis and test if fluxes were associated with sediment chemistry, including percent organic matter in sediment (%OM) and sediment polyphenolic concentrations (polyphenols), and microbial communities, including abundance (abund), diversity (divers), and composition (comp) of methanogens (*mcrA*) and methanotrophs encoding *mmoX* or *pmoA* genes. All models included climate, including water temperature (temp), and other environmental variables, e.g. dissolved organic carbon concentrations (DOC) and organic matter quality in porewater measured by specific ultraviolet absorbance (SUVA). Models were compared with the Akaike information criterion with R^2^ value from best supported model (Table S2). Shading of boxes and text correspond to model support with darker colours indicating greater support (Table S2). Arrows point at modeled variables, with their sizes proportional to the mean effect of one variable on another estimated by path analysis and colours corresponding with direction of effect. Effect sizes plotted for predictors included in all models (i.e. climate and environment) were from the best supported model. Correlations among abund variables omitted for clarity.

**Figure 4.**
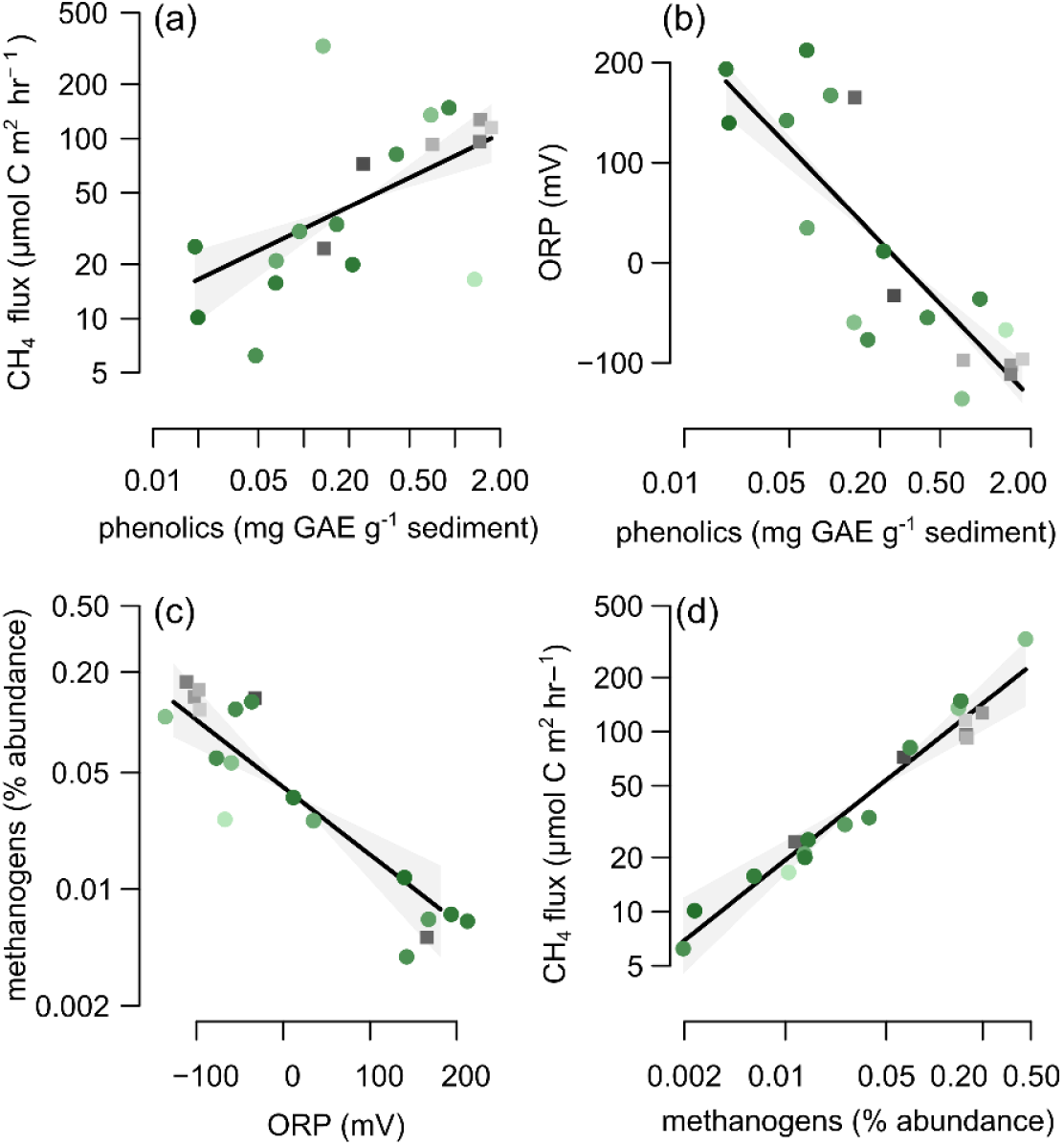
Drivers of littoral methane gas emissions. Diffusive CH_4_ fluxes per m^2^ of sediment increased with (**a**) polyphenolic concentrations expressed in gallic acid equivalents (GAE) per g dry weight of sediment. This association was predicted to arise in the best supported path analysis (Fig. 3, Table S2), because (**b**) higher polyphenolic concentrations reduced oxidation reduction potential (ORP), and **(c)** lower ORP increased the summed relative abundance of methanogens that were present in >50% of samples and were individually and positively correlated with CH_4_ fluxes (Fig. 2c). **(d)** CH_4_ fluxes then proximally increased with the summed relative abundance of these methanogens. Points are site-level model predictions with fit (R^2^) reported in Fig. 3 and coloured as in Fig. 1, indicating either percent of grassland in surrounding catchments for littoral deposition zones (circles) or latitude for macrophyte beds (squares). Lines are estimated model fits ± 95% CI from path analyses that simultaneously account for other effects shown in Fig. 3. For (**a**), we estimated the total effect of polyphenolics on CH_4_, that is, the sum of the direct and indirect effects weighted by AICc-estimated model support (Table S2).

There was no evidence that other organic matter loaded into sediments alongside the polyphenolics was more important for predicting GHG emissions. Under this scenario, organic matter itself would have been a better indirect predictor of CH_4_ mediated via redox potential and subsequently methanogens, or a better direct predictor of CH_4_ than the polyphenolics. However, replacing polyphenolic concentrations by organic matter content increased AICc in *P2* and *P3* by 11.5 and 1.3, respectively, indicating that these were not better supported models. Likewise, using a mathematical model of geochemical reactions, we found that the dissolution of two model polyphenolics into lake water reduced redox potentials to levels consistent with our observations and more so than other compounds (Fig. S2). Polyphenolics themselves accumulated with more OM and as SUVA increased from 0.2 to 6.9 L mg C^-1^ m^-1^, likely reflecting a greater contribution of terrestrial sources (Fig. 3). Polyphenolics may have also accumulated in sediment because lower redox potentials created anaerobic conditions that better preserved complex organic substrates, but we found little support for this pathway. AICc increased by 27.3 and 24.3 when we swapped the directionality between redox potential and sediment polyphenolic concentrations in *P1* and *P2*, respectively.

Although all copy numbers of the methanogen marker *mcrA* were predicted to increase diffusive CH_4_ emissions directly, such as by enhancing total methanogen activity, and by increasing methanogen diversity, our sites offered negligible support for models *P4* and *P5*, respectively (AIC weight = 0.2% for both). We similarly found little evidence that (*P6*) methanotroph abundances helped to explain variation in CH_4_ emissions (0.1% support), and there was no correlation between CH_4_ and CO_2_ after accounting for model predictors (partial correlation, r = 0.28, p = 0.134). Copy numbers for the methanotroph marker *pmoA* but not *mmoX* did, however, increased under higher sediment OM concentrations, and were subsequently associated with CO_2_ fluxes changing from a mean sink of –1.5 (95% CI: –10.7 to 4.5) mmol C hr^-1^ to mean source of 27.0 (17.0 to 42.2) mmol C hr^-1^ (Fig. 5). None of the other environmental predictors (i.e. pH, DOC concentration, temperature, latitude) had statistically significant effects except for temperature on CO_2_ concentrations and the models generally fitted the data well (R^2^ for individual responses = 0.23 to 0.85, Fig. 3).

**Figure 5.**
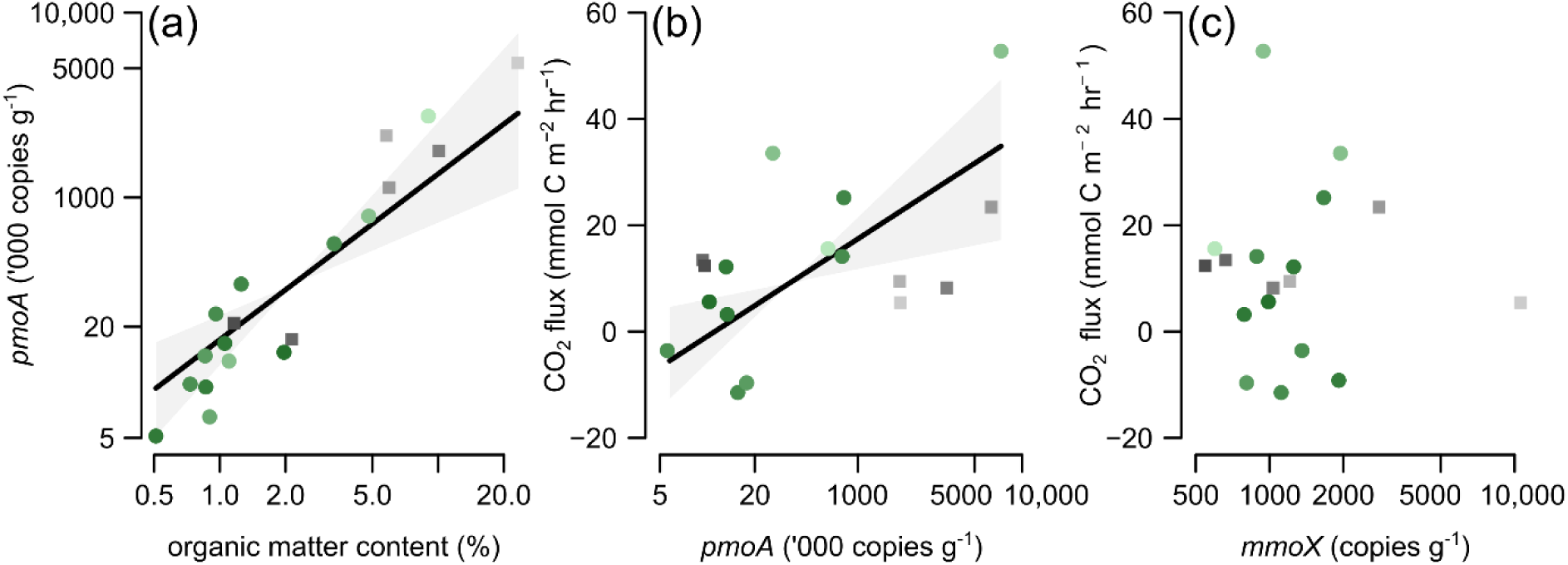
Microbial drivers of littoral carbon dioxide emissions. **(a)** Abundances of bacteria containing the particulate methane monooxygenase (*pmoA*) gene (thousands of copies per g dry weight of sediment) increased with the percentage of organic matter in sediment. CO_2_ fluxes per m^2^ of sediment subsequently (b) increased with the abundance of *pmoA* and but showed no association with the abundance of methane methylocella (*mmoX*), which is another methane-oxidising marker. Points are site-level predictions with fit (R^2^) reported in Fig. 3 and coloured as in Fig. 1, indicating either percent of grassland in surrounding catchments for littoral deposition zones (circles) or latitude for macrophyte beds (squares). Lines are estimated model fits ± 95% CI from the best supported path analysis that simultaneously account for other effects shown in Fig. 3.

## Discussion

Our study helps to resolve conflicting evidence about the role of sediment chemistry in controlling aquatic methane fluxes. In contrast to lab studies leaching plant litterfall or field-based studies of peatlands (Emilson and others 2018; Fenner and Freeman 2020), our results suggest polyphenolics neither (*P3*) inactivated enzymes involved in decomposition of organic matter nor strongly (*P4*) inhibited abundances of methanogens. Instead, our results suggest polyphenolics favour methanogenesis by reducing redox potentials to levels that promote certain methanogen taxa (*P2*) rather than by directly changing the electrochemical favourability of reactions (*P1*) or methanogen abundance (*P4*) and diversity (*P5*). A key difference with our study is that polyphenolic concentrations were lower compared to others (Emilson and others 2018; Fenner and Freeman 2020), which we attribute to the surrounding catchments that overwhelmingly supply OM to littoral deposition zones. Our study also provides new evidence from littoral sediments indicating that lake CH_4_ emissions depend on abundances of methanotrophic organisms, as predicted by (*P6*). We now advance past work by demonstrating that these effects of methanotrophs are relatively minor in shallow sediments as compared with other drivers of littoral methane fluxes.

Our study catchments were dominated by grasslands that have lower foliar polyphenolic concentrations (Alonso and others 2001) than woodland, macrophyte, or peat-forming vegetation (Emilson and others 2018; Zak and others 2019). Polyphenolic concentrations in depositional zones correspondingly decreased by an estimated 16-times (95% CI: 1.2 to 196.7) as grassland cover increased over its range of values in our surrounding catchments (linear model R^2^ = 0.28; Fig. 1). Most polyphenolics are also water soluble (Min and others 2015), so can be quickly leached away in lakes unlike in controlled microcosms or peatlands with longer water residence time. Lake sediments may therefore have lower polyphenolic concentrations than expected from their litter inputs. The only two studies, to our knowledge, which have also measured polyphenolics in lake sediments have found similarly low concentrations that are several tens-of-times lower than in surrounding vegetation (Quayyum and others 1999; Cieślewicz 2014). These studies suggest our results are likely representative of lake sediments elsewhere. Furthermore, the contrasting chemistry between macrophyte– and inflow-dominated sites, and large range of catchment composition and thus OM inputs that we sampled, further suggest our findings are generally representative.

The mixed responses of methane fluxes to polyphenolic concentrations reported previously may also have arisen because of variation in microbial community composition. Studies reporting a negative correlation between CH_4_ and polyphenolic concentrations in lake sediment have reported tens-of-times higher polyphenolic concentrations than our field data but much lower CH_4_ production (Emilson and others 2018; Praetzel and others 2020 report only CH_4_ in comparable units). One explanation for this result is high polyphenolic concentrations limit some pathways of biological CH_4_ production. The study reporting a negative correlation between CH_4_ and polyphenolic concentrations was dominated by Methanobacteriaceae that typically use hydrogenotrophic methanogenesis (Yakimovich and others 2018).

Hydrogenotrophic methanogens require CO_2_, of which production can be inhibited by reduced conditions associated with high polyphenolic concentrations (Freeman and others 2001; Wilmoth and others 2021). Recent evidence has instead shown that acetoclastic and methylotrophic methanogens can breakdown a model polyphenol in anoxic soils (McGivern and others 2021). Amendments of phenol-rich OM to soils elsewhere increased CH_4_ production only where acetoclastic methanogenesis was important (Medvedeff and others 2015). As our study sites were dominated by taxa that use both acetoclastic and methylotrophic pathways (Liu and Whitman 2008), methanogens could have remained active in our sites, and a higher abundance or diversity of methanogens would not necessarily translate into greater activity. Instead, shifts towards lower redox may have increased the abundance of key methanogens (Conrad and others 1987; Zhuang and others 2018). Because lake sediments are generally dominated by Methanosarcinales and Methanomicrobiales (Wen and others 2017), this mechanism may generalise to many lakes and highlights the importance of improving the identification of uncultivated microbes like the many detected here.

A limitation of our study was that we did not directly compare the effect of polyphenolics with many other compounds that can influence lake methane emissions, such as metals (e.g. iron), hydrides (e.g. H_2_S), and organic decomposition products (e.g. acetate). Geochemical modelling indicates that only polyphenolics could reduce redox potential to levels that favour methanogenesis (Fig. S2), though this approach could not capture the full mixture of compounds that create similar redox potentials observed *in situ*. Other measures of OM (e.g. DOC concentrations and OM complexity) were also still poorer predictors of our field data than polyphenolics alone. Although the presence of Fe(II) is thought to help reduce redox potential and facilitate hydrogenotrophic methanogenesis (Jiang and others 2013), we found few methanogen taxa characteristic of hydrogenotrophic pathways in our study sites, suggesting that this process is unlikely to have been important. A less likely explanation for our results is that polyphenolics better represent the reactivity of OM than bulk pools without necessarily being the causal drivers. However, studies that experimentally added polyphenolics to soils have found that they can be reactive and quickly degraded by diverse microorganisms (Steinmetz and others 2019; McGivern and others 2021).

We also aimed to test a specific hypothesis about the importance of polyphenolics for methanogenesis rather than fully disentangle all potential mechanisms of sediment methane production. Substrate competition, such as from more energetically favourable metabolic pathways involving sulfate-or nitrate-reducing bacteria (Kuivila and others 1989; Percheron and others 1999), and other sediment characteristics, e.g. mineralogy (Xiao and others 2022), may contribute towards some of the unexplained variation in our results. Geological weathering of iron oxides by some archaea can also be an important predictor of lake CH_4_ emissions by catalysing anaerobic oxidation of methane (Ettwig and others 2016), but we again found little evidence of ANMEs in our study sites. Finally, we did not measure overlying water chemistry. Past work in littoral sediments suggests that surface water has a smaller effect on CH_4_ concentrations than porewater chemistry, and this effect is likely mediated through shifts in microbial composition rather than direct chemical effects, such as nutrient concentrations (Tanentzap and others 2019).

Ecosystem models predict methane emissions with large uncertainty, even when microbial population dynamics are considered (Chadburn and others 2020). Our study suggests that this uncertainty could be reduced by modelling the chemical composition of OM inputs to freshwaters from surrounding vegetation. Specifically, electrochemical models can be used to predict redox potential from polyphenolic concentrations of different OM sources (e.g. Novak Jovanović and Miličević 2017). Predicted redox potential could then inform microbial dynamics and model-estimated methane fluxes. As vegetation in and around lakes shift with a changing climate, knowledge of how their chemistry impacts methane emissions can ultimately aid climate mitigation efforts.

## Acknowledgements

This work was funded by a Royal Society Research Grant #180279 to A.J.T.

## Data availability

Data, metadata, and R code used to generate the results of this study are available in Zenodo (https://doi.org/10.5281/zenodo.14728117). Sequence data have been deposited in the NCBI Sequence Read Archive (accession numbers in Table S1)

## Author contributions

AJT designed the research, AJT and SGW undertook the field work, AJT, OK, and YZ undertook the lab work, OK developed new sequencing methods, SGW performed geospatial analyses, AJT analysed the data and wrote the manuscript with input from all authors.

## References

1. Aben RCH, Barros N, van Donk E, Frenken T, Hilt S, Kazanjian G, Lamers LPM, Peeters ETHM, Roelofs JGM, de Senerpont Domis LN, Stephan S, Velthuis M, Van de Waal DB, Wik M, Thornton BF, Wilkinson J, DelSontro T, Kosten S. 2017. Cross continental increase in methane ebullition under climate change. Nature Communications 8:1682.

2. Ainsworth EA, Gillespie KM. 2007. Estimation of total phenolic content and other oxidation substrates in plant tissues using Folin–Ciocalteu reagent. Nature Protocols 2:875–7.

3. Alonso I, Hartley S e., Thurlow M. 2001. Competition between heather and grasses on Scottish moorlands: Interacting effects of nutrient enrichment and grazing regime. Journal of Vegetation Science 12:249–60.

4. Angel R, Claus P, Conrad R. 2012. Methanogenic archaea are globally ubiquitous in aerated soils and become active under wet anoxic conditions. ISME J 6:847–62.

5. Barbosa PM, Melack JM, Amaral JHF, Linkhorst A, Forsberg BR. 2021. Large seasonal and habitat differences in methane ebullition on the Amazon floodplain. Journal of Geophysical Research: Biogeosciences 126:e2020JG005911.

6. Bastviken D, Cole JJ, Pace ML, Van de Bogert MC. 2008. Fates of methane from different lake habitats: Connecting whole-lake budgets and CH4 emissions. Journal of Geophysical Research: Biogeosciences 113:G02024.

7. Bertolet BL, West WE, Armitage DW, Jones SE. 2019. Organic matter supply and bacterial community composition predict methanogenesis rates in temperate lake sediments. Limnology and Oceanography Letters 4:164–72.

8. Bodmer, P., Vroom, R. J. E., Stepina, T., del Giorgio, P. A., & Kosten, S. (2024). Methane dynamics in vegetated habitats in inland waters: quantification, regulation, and global significance. Frontiers in Water 5:1332968.

9. Chadburn SE, Aalto T, Aurela M, Baldocchi D, Biasi C, Boike J, Burke EJ, Comyn-Platt E, Dolman AJ, Duran-Rojas C, Fan Y, Friborg T, Gao Y, Gedney N, Göckede M, Hayman GD, Holl D, Hugelius G, Kutzbach L, Lee H, Lohila A, Parmentier F-JW, Sachs T, Shurpali NJ, Westermann S. 2020. Modeled microbial dynamics explain the apparent temperature sensitivity of wetland methane emissions. Global Biogeochemical Cycles 34:e2020GB006678.

10. Chong I-G, Jun C-H. 2005. Performance of some variable selection methods when multicollinearity is present. Chemometrics and Intelligent Laboratory Systems 78:103– 12.

11. Chowdhury TR, Dick RP. 2013. Ecology of aerobic methanotrophs in controlling methane fluxes from wetlands. Applied Soil Ecology 65:8–22.

12. Cieślewicz J. 2014. Polyphenolic compounds in lacustrine sediments. Polish Journal of Environmental Studies 23:1965–73.

13. Conrad R, Lupton FS, Zeikus JG 1987. Hydrogen metabolism and sulfate-dependent inhibition of methanogenesis in a eutrophic lake sediment (Lake Mendota). FEMS Microbiology Ecology 3:107–115.

14. Deemer BR, Holgerson MA. 2021. Drivers of methane flux differ between lakes and reservoirs, complicating global upscaling efforts. Journal of Geophysical Research: Biogeosciences 126:e2019JG005600.

15. DelSontro T, Boutet L, St-Pierre A, del Giorgio PA, Prairie YT. 2016. Methane ebullition and diffusion from northern ponds and lakes regulated by the interaction between temperature and system productivity. Limnology and Oceanography 61:S62–77.

16. Delwiche KB, Knox SH, Malhotra A, Fluet-Chouinard E, McNicol G, Feron S, Ouyang Z, Papale D, Trotta C, Canfora E, Cheah Y-W, Christianson D, Alberto MCR, Alekseychik P, Aurela M, Baldocchi D, Bansal S, Billesbach DP, Bohrer G, Bracho R, Buchmann N, Campbell DI, Celis G, Chen J, Chen W, Chu H, Dalmagro HJ, Dengel S, Desai AR, Detto M, Dolman H, Eichelmann E, Euskirchen E, Famulari D, Fuchs K, Goeckede M, Gogo S, Gondwe MJ, Goodrich JP, Gottschalk P, Graham SL, Heimann M, Helbig M, Helfter C, Hemes KS, Hirano T, Hollinger D, Hörtnagl L, Iwata H, Jacotot A, Jurasinski G, Kang M, Kasak K, King J, Klatt J, Koebsch F, Krauss KW, Lai DYF, Lohila A, Mammarella I, Belelli Marchesini L, Manca G, Matthes JH, Maximov T, Merbold L, Mitra B, Morin TH, Nemitz E, Nilsson MB, Niu S, Oechel WC, Oikawa PY, Ono K, Peichl M, Peltola O, Reba ML, Richardson AD, Riley W, Runkle BRK, Ryu Y, Sachs T, Sakabe A, Sanchez CR, Schuur EA, Schäfer KVR, Sonnentag O, Sparks JP, Stuart-Haëntjens E, Sturtevant C, Sullivan RC, Szutu DJ, Thom JE, Torn MS, Tuittila E-S, Turner J, Ueyama M, Valach AC, Vargas R, and others. 2021. FLUXNET-CH_4_: a global, multi-ecosystem dataset and analysis of methane seasonality from freshwater wetlands. Earth System Science Data 13:3607–89.

17. Desrosiers K, DelSontro T, del Giorgio PA. 2022. Disproportionate contribution of vegetated habitats to the CH4 and CO2 budgets of a boreal lake. Ecosystems 25:1522–41.

18. Ding H, Hu Q, Cai M, Cao C, Jiang Y. 2022. Effect of dissolved organic matter (DOM) on greenhouse gas emissions in rice varieties. Agriculture, Ecosystems & Environment 330:107870.

19. Ellenbogen JB, Borton MA, McGivern BB, Cronin DR, Hoyt DW, Freire-Zapata V, McCalley CK, Varner RK, Crill PM, Wehr RA, Chanton JP, Woodcroft BJ, Tfaily MM, Tyson GW, Rich VI, Wrighton KC. 2023. Methylotrophy in the mire: direct and indirect routes for methane production in thawing permafrost. mSystems 9:e00698–23.

20. Emerson JB, Varner RK, Wik M, Parks DH, Neumann RB, Johnson JE, Singleton CM, Woodcroft BJ, Tollerson R, Owusu-Dommey A, Binder M, Freitas NL, Crill PM, Saleska SR, Tyson GW, Rich VI. 2021. Diverse sediment microbiota shape methane emission temperature sensitivity in Arctic lakes. Nature Communications 12:5815.

21. Emilson EJS, Carson MA, Yakimovich KM, Osterholz H, Dittmar T, Gunn JM, Mykytczuk NCS, Basiliko N, Tanentzap AJ. 2018. Climate-driven shifts in sediment chemistry enhance methane production in northern lakes. Nature Communications 9:1801.

22. Erkkilä K-M, Ojala A, Bastviken D, Biermann T, Heiskanen JJ, Lindroth A, Peltola O, Rantakari M, Vesala T, Mammarella I. 2018. Methane and carbon dioxide fluxes over a lake: comparison between eddy covariance, floating chambers and boundary layer method. Biogeosciences 15:429–45.

23. Fenner N, Freeman C. 2020. Woody litter protects peat carbon stocks during drought. Nature Climate Change 10:363–9.

24. Freeman C, Ostle N, Kang H. 2001. An enzymic ‘latch’ on a global carbon store. Nature 409:149–149.

25. Gonzalez-Valencia R, Magana-Rodriguez F, Gerardo-Nieto O, Sepulveda-Jauregui A, Martinez-Cruz K, Walter Anthony K, Baer D, Thalasso F. 2014. In situ measurement of dissolved methane and carbon dioxide in freshwater ecosystems by off-axis integrated cavity output spectroscopy. Environmental Science & Technology 48:11421–8.

26. Goodrich JP, Varner RK, Frolking S, Duncan BN, Crill PM. 2011. High-frequency measurements of methane ebullition over a growing season at a temperate peatland site. Geophysical Research Letters 38:L07404.

27. Grace JB, Irvine KM. 2020. Scientist’s guide to developing explanatory statistical models using causal analysis principles. Ecology 101:e02962.

28. Grasset C, Abril G, Guillard L, Delolme C, Bornette G. 2016. Carbon emission along a eutrophication gradient in temperate riverine wetlands: effect of primary productivity and plant community composition. Freshwater Biology 61:1405–20.

29. Grasset C, Mendonça R, Villamor Saucedo G, Bastviken D, Roland F, Sobek S. 2018. Large but variable methane production in anoxic freshwater sediment upon addition of allochthonous and autochthonous organic matter. Limnology and Oceanography 63:1488–501.

30. Heslop J, Walter Anthony K, Zhang M. 2017. Utilizing pyrolysis GC-MS to characterize organic matter quality in relation to methane production in a thermokarst lake sediment core. Organic Geochemistry 103:43–50.

31. Hodgkins SB, Tfaily MM, McCalley CK, Logan TA, Crill PM, Saleska SR, Rich VI, Chanton JP. 2014. Changes in peat chemistry associated with permafrost thaw increase greenhouse gas production. Proceedings of the National Academy of Sciences 111:5819– 24.

32. Holgerson MA, Raymond PA. 2016. Large contribution to inland water CO2 and CH4 emissions from very small ponds. Nature Geoscience 9:222–6.

33. Jin Q, Kirk MF. 2018. pH as a primary control in environmental microbiology: 1. thermodynamic perspective. Frontiers in Environment Science 6:21.

34. Juutinen S, Alm J, Larmola T, Huttunen JT, Morero M, Martikainen PJ, Silvola J. 2003. Major implication of the littoral zone for methane release from boreal lakes. Global Biogeochemical Cycles 17:1117.

35. Kane ES, Veverica TJ, Tfaily MM, Lilleskov EA, Meingast KM, Kolka RK, Daniels AL, Chimner RA. 2019. Reduction-oxidation potential and dissolved organic matter composition in northern peat soil: interactive controls of water table position and plant functional groups. Journal of Geophysical Research: Biogeosciences 124:3600–17.

36. Kuhn MA, Thompson LM, Winder JC, Braga LPP, Tanentzap AJ, Bastviken D, Olefeldt D. 2021. Opposing effects of climate and permafrost thaw on CH4 and CO2 emissions from northern lakes. AGU Advances 2:e2021AV000515.

37. Kuivila KM, Murray JW, Devol AH, Novelli PC. 1989. Methane production, sulfate reduction and competition for substrates in the sediments of Lake Washington. Geochimica et Cosmochimica Acta 53:409–16.

38. LaRowe DE, Van Cappellen P. 2011. Degradation of natural organic matter: A thermodynamic analysis. Geochimica et Cosmochimica Acta 75:2030–42.

39. Lavonen EE, Kothawala DN, Tranvik LJ, Gonsior M, Schmitt-Kopplin P, Köhler SJ. 2015. Tracking changes in the optical properties and molecular composition of dissolved organic matter during drinking water production. Water Research 85:286–94.

40. Lefcheck JS. 2016. piecewiseSEM: Piecewise structural equation modelling in r for ecology, evolution, and systematics. Methods in Ecology and Evolution 7:573–9.

41. Lew S, Glińska-Lewczuk K. 2018. Environmental controls on the abundance of methanotrophs and methanogens in peat bog lakes. Science of The Total Environment 645:1201–11.

42. Li H. 2018. Minimap2: pairwise alignment for nucleotide sequences. Bioinformatics 34:3094– 100.

43. Liu Y, Whitman WB. 2008. Metabolic, phylogenetic, and ecological diversity of the methanogenic Archaea. Annals of the New York Academy of Sciences 1125:171–89.

44. Marotta H, Pinho L, Gudasz C, Bastviken D, Tranvik LJ, Enrich-Prast A. 2014. Greenhouse gas production in low-latitude lake sediments responds strongly to warming. Nature Climate Change 4:467–70.

45. McGinnis DF, Greinert J, Artemov Y, Beaubien SE, Wüest A. 2006. Fate of rising methane bubbles in stratified waters: How much methane reaches the atmosphere? Journal of Geophysical Research: Oceans 111: C09007.

46. McGivern BB, Tfaily MM, Borton MA, Kosina SM, Daly RA, Nicora CD, Purvine SO, Wong AR, Lipton MS, Hoyt DW, Northen TR, Hagerman AE, Wrighton KC. 2021. Decrypting bacterial polyphenol metabolism in an anoxic wetland soil. Nature Communications 12:2466.

47. McMurdie PJ, Holmes S. 2014. Waste not, want not: why rarefying microbiome data is inadmissible. PLOS Computational Biology 10:e1003531.

48. Medvedeff CA, Bridgham SD, Pfeifer-Meister L, Keller JK. 2015. Can Sphagnum leachate chemistry explain differences in anaerobic decomposition in peatlands? Soil Biology and Biochemistry 86:34–41.

49. Mevik B-H, Wehrens R. 2007. The pls package: principal component and partial least squares regression in R. Journal of Statistical Software 18:1–23.

50. Min BR, Solaiman S, Waldrip HM, Parker D, Todd RW, Brauer D. 2020. Dietary mitigation of enteric methane emissions from ruminants: A review of plant tannin mitigation options. Animal Nutrition 6:231–46.

51. Muggeo VMR 2017. Interval estimation for the breakpoint in segmented regression: a smoothed score-based approach. Australian & New Zealand Journal of Statistics 59:311–322.

52. Murphy MV 2022. semEff: Automatic calculation of effects for piecewise structural equation models. Retrieved from https://CRAN.R-project.org/package=semEff.

53. Novak Jovanović I, Miličević A. 2017. A model for the estimation of oxidation potentials of polyphenols. Journal of Molecular Liquids 241:255–9.

54. Oksanen J, Simpson G, Blanchet FG, Kindt R, Legendre P, Minchin P, and others. 2022. vegan: community ecology package. Retrieved from https://cran.r-project.org/package=vegan

55. Peacock M, Audet J, Bastviken D, Cook S, Evans CD, Grinham A, Holgerson MA, Högbom L, Pickard AE, Zieliński P, Futter MN. 2021. Small artificial waterbodies are widespread and persistent emitters of methane and carbon dioxide. Global Change Biology 27:5109– 23.

56. Percheron G, Bernet N, Moletta R. 1999. Interactions between methanogenic and nitrate reducing bacteria during the anaerobic digestion of an industrial sulfate rich wastewater. FEMS Microbiology Ecology 29:341–50.

57. Praetzel LSE, Plenter N, Schilling S, Schmiedeskamp M, Broll G, Knorr K-H. 2020. Organic matter and sediment properties determine in-lake variability of sediment CO_2_ and CH_4_ production and emissions of a small and shallow lake. Biogeosciences 17:5057–78.

58. Putkinen A, Tuittila E-S, Siljanen HMP, Bodrossy L, Fritze H. 2018. Recovery of methane turnover and the associated microbial communities in restored cutover peatlands is strongly linked with increasing Sphagnum abundance. Soil Biology and Biochemistry 116:110–9.

59. Quayyum HA, Mallik AU, Lee PF. 1999. Allelopathic potential of aquatic plants associated with wild rice (*Zizania palustris*): I. Bioassay with plant and lake sediment samples. Journal of Chemical Ecology 25:209–20.

60. R Core Team 2022. R: A language and environment for statistical computing. R Foundation for Statistical Computing. https://www.R-project.org/.

61. Rahman MT, Crombie A, Chen Y, Stralis-Pavese N, Bodrossy L, Meir P, McNamara NP, Murrell JC. 2011. Environmental distribution and abundance of the facultative methanotroph *Methylocella*. ISME J 5:1061–6.

62. Rodríguez-Pérez H, Ciuffreda L, Flores C. 2021. NanoCLUST: a species-level analysis of 16S rRNA nanopore sequencing data. Bioinformatics 37:1600–1.

63. Rosentreter JA, Borges AV, Deemer BR, Holgerson MA, Liu S, Song C, Melack J, Raymond PA, Duarte CM, Allen GH, Olefeldt D, Poulter B, Battin TI, Eyre BD. 2021. Half of global methane emissions come from highly variable aquatic ecosystem sources. Nature Geoscience 14:225–30.

64. Schloss PD. 2024. Rarefaction is currently the best approach to control for uneven sequencing effort in amplicon sequence analyses. mSphere 9:e00354–23.

65. Seeberg-Elverfeldt J, Schlüter M, Feseker T, Kölling M. 2005. Rhizon sampling of porewaters near the sediment-water interface of aquatic systems. Limnology and Oceanography: Methods 3:361–71.

66. Sereika M, Kirkegaard RH, Karst SM, Michaelsen TY, Sørensen EA, Wollenberg RD, Albertsen M. 2022. Oxford Nanopore R10.4 long-read sequencing enables the generation of near-finished bacterial genomes from pure cultures and metagenomes without short-read or reference polishing. Nature Methods 19:823–6.

67. Sieczko AK, Duc NT, Schenk J, Pajala G, Rudberg D, Sawakuchi HO, Bastviken D. 2020. Diel variability of methane emissions from lakes. Proceedings of the National Academy of Sciences 117:21488–94.

68. Steinmetz Z, Kurtz MP, Zubrod JP, Meyer AH, Elsner M, Schaumann GE. 2019. Biodegradation and photooxidation of phenolic compounds in soil—A compound-specific stable isotope approach. Chemosphere 230:210–8.

69. Tanentzap AJ, Fitch A, Orland C, Emilson EJS, Yakimovich KM, Osterholz H, Dittmar T. 2019. Chemical and microbial diversity covary in fresh water to influence ecosystem functioning. Proceedings of the National Academy of Sciences116:24689–95.

70. Walpen N, Getzinger GJ, Schroth MH, Sander M. 2018. Electron-donating phenolic and electron-accepting quinone moieties in peat dissolved organic matter: quantities and redox transformations in the context of peat biogeochemistry. Environmental Science & Technology 52:5236–45.

71. West WE, Coloso JJ, Jones SE. 2012. Effects of algal and terrestrial carbon on methane production rates and methanogen community structure in a temperate lake sediment. Freshwater Biology 57:949–55.

72. West WE, Creamer KP, Jones SE. 2016. Productivity and depth regulate lake contributions to atmospheric methane. Limnology and Oceanography 61:S51–61.

73. Wetzel RG. 2001. Limnology: Lake and River Ecosystems. San Diego: Academic Press

74. Wilmoth JL, Schaefer JK, Schlesinger DR, Roth SW, Hatcher PG, Shoemaker JK, Zhang X. 2021. The role of oxygen in stimulating methane production in wetlands. Global Change Biology 27:5831–47.

75. Winder JC, Braga LPP, Kuhn MA, Thompson LM, Olefeldt D, Tanentzap AJ. 2023. Climate warming has direct and indirect effects on microbes associated with carbon cycling in northern lakes. Global Change Biology 29:3039–53.

76. Xiao K-Q, Moore OW, Babakhani P, Curti L, Peacock CL. 2022. Mineralogical control on methylotrophic methanogenesis and implications for cryptic methane cycling in marine surface sediment. Nature Communications 13:2722.

77. Xie F, Zhao S, Zhan X, Zhou Y, Li Y, Zhu W, Pope PB, Attwood GT, Jin W, Mao S. 2024. Unraveling the phylogenomic diversity of Methanomassiliicoccales and implications for mitigating ruminant methane emissions. Genome Biology 25:32.

78. Yakimovich KM, Emilson EJS, Carson MA, Tanentzap AJ, Basiliko N, Mykytczuk NCS. 2018. Plant litter type dictates microbial communities responsible for greenhouse gas production in amended lake sediments. Frontiers in Microbiology 9:2662.

79. Yakimovich KM, Orland C, Emilson EJS, Tanentzap AJ, Basiliko N, Mykytczuk NCS. 2020. Lake characteristics influence how methanogens in littoral sediments respond to terrestrial litter inputs. The ISME Journal 14:2153–63.

80. Yang S, Liebner S, Alawi M, Ebenhöh O, Wagner D. 2014. Supplement to: Taxonomic database and cut-off value for processing mcrA gene 454 pyrosequencing data by MOTHUR. 10.5880/GFZ.4.5.2014.001. Last accessed 13/03/2023

81. Yavitt JB, Williams CJ, Wieder RK. 1997. Production of methane and carbon dioxide in peatland ecosystems across North America: Effects of temperature, aeration, and organic chemistry of peat. Geomicrobiology Journal 14:299–316.

82. Yvon-Durocher G, Allen AP, Bastviken D, Conrad R, Gudasz C, St-Pierre A, Thanh-Duc N, del Giorgio PA. 2014. Methane fluxes show consistent temperature dependence across microbial to ecosystem scales. Nature 507:488–91.

83. Zak D, Roth C, Unger V, Goldhammer T, Fenner N, Freeman C, Jurasinski G. 2019. Unraveling the importance of polyphenols for microbial carbon mineralization in rewetted riparian peatlands. Frontiers in Environmental Science 7:147.

84. Zhuang G-C, Montgomery A, Sibert RJ, Rogener M-K, Samarkin VA, Joye SB. 2018. Effects of pressure, methane concentration, sulfate reduction activity, and temperature on methane production in surface sediments of the Gulf of Mexico. Limnology and Oceanography 63:2080–2092.

