## Supplement Materials for "Sediment chemistry controls methane emissions from lake littoral zones"

**Supplementary Methods S1 – Additional methods**

*Lake characteristics and land cover classification*

We first delineated catchments with the 25 m European digital elevation model (DEM), v1.1 (European Environment Agency 2016) in R v4.0 using whitebox v2.1.0 (Lindsay 2016). Prior to delineation, single cell pits and sinks were filled using breach depression. In four sites (Allt Ceann, Lochy, Lubnaig, and Tay), accurate catchment delineation was difficult at the resolution of DEM because the sampled inflows were relatively small (<3 m wide). We therefore used the EU-Hydro river network database v1.3 (European Environment Agency 2020) and OpenStreetMap (<https://www.openstreetmap.org/>) river vectors to burn the channel into the DEM. This approach sets the elevation of pixels along the river vector to lower values than their surroundings to improve the recognition of known stream locations in subsequent processing steps. Burning in known stream locations is particularly useful for delineating small streams as their channels are difficult to detect at lower DEM resolutions (Goulde, 2014). In these smaller streams we also used HydroLakes polygons (Messager and others 2016) to flatten the lake area to reduce errors associated with flat lake bends.

After all catchments were delineated (total area: 8 to 19,521 ha), they were intersected with a 20 m resolution land cover map of the entire UK in 2018 (Morton and others 2020). All water pixels were removed and we summed the total of land area in each catchment under grassland (acid or improved), heather, heather grassland, bog, and arable land cover.

We used HydroLakes to extract area and depth for lakes ≥0.2 km^2^ (Messager and others 2016), with values for the two sites <0.2 km^2^ derived from the EU-Hydro river network database (European Environment Agency 2020).

*Modifications to qPCR protocols*

We used the following settings for the qPCRs described in the Main Text:

*mmoX*: initial denaturation at 95°C for 10 min, 45 cycles of 95°C for 15 s, 68°C for 1 min, 72°C for 1 min, and a final extension at 72°C for 10 min

*pmoA* type Ia: initial denaturation at 95°C for 10 min, 42 cycles of 95°C for 30 s, 54 °C for 35 s, 72°C for 45 s, and a final extension at 72°C for 10 min

*pmoA* type Ib: same as type Ia except for an annealing step of 64°C for 45 s

*pmoA* type II: touchdown PCR was performed with denaturation at 95°C for 10 min, 12 cycles of 95°C for 15 s, 63°C to 52°C (decreased by 1°C for each cycle) for 45 s, 72°C for 30 s, followed by 30 cycles of 95°C for 15 s, 52°C for 45 s, 72°C for 30 s and a final extension at 72°C for 10 min.

**Supplementary References**

Ellenbogen JB, Borton MA, McGivern BB, Cronin DR, Hoyt DW, Freire-Zapata V, McCalley CK, Varner RK, Crill PM, Wehr RA, Chanton JP, Woodcroft BJ, Tfaily MM, Tyson GW, Rich VI, Wrighton KC. 2023. Methylotrophy in the mire: direct and indirect routes for methane production in thawing permafrost. mSystems 9:e00698-23.

European Environment Agency. 2016. European Digital Elevation Model (EU-DEM), version 1.1. <http://land.copernicus.eu/pan-european/satellite-derived-products/eu-dem/eu-dem-v1.1> [accessed 2 May 2022]

European Environment Agency. 2020. EU-Hydro – River Network Database, version 1.3. <https://land.copernicus.eu/imagery-in-situ/eu-hydro/eu-hydro-river-network-database> [accessed 2 May 2022]

Goulden T, Hopkinson C, Jamieson R, Sterling S. 2014. Sensitivity of watershed attributes to spatial resolution and interpolation method of LiDAR DEMs in three distinct landscapes. Water Resources Research 50:1908–1927.

Lindsay JB. 2016. Whitebox GAT: A case study in geomorphometric analysis. Computers & Geosciences 95:75–84.

Liu Y, Whitman WB. 2008. Metabolic, phylogenetic, and ecological diversity of the methanogenic Archaea. Annals of the New York Academy of Sciences 1125:171–89.

Livingstone DA. 1963. Data of geochemistry: chemical composition of rivers and lakes. U.S. Geological Survey, Washington D.C., p. 64. https://doi.org/10.3133/pp440G

Messager ML, **Lehner B**, Grill G, Nedeva I, Schmitt O. 2016. Estimating the volume and age of water stored in global lakes using a geo-statistical approach. Nature Communications 7:13603 doi: 10.1038/ncomms13603

Morton RD, Marston CG, O’Neil AW, Rowland CS. 2020. Land Cover Map 2018 (20m classified pixels, GB). NERC Environmental Information Data Centre. doi: 10.5285/b3dfc4c7-c9bd-4a02-bed8-46b2a41be04a

Ou Y-F, Dong H-P, McIlroy SJ, Crowe SA, Hallam SJ, Han P, Kallmeyer J, Simister RL, Vuillemin A, Leu AO, Liu Z, Zheng Y-L, Sun Q-L, Liu M, Tyson GW, Hou L-J. 2022. Expanding the phylogenetic distribution of cytochrome b-containing methanogenic archaea sheds light on the evolution of methanogenesis. ISME J 16:2373–87.

Parkhurst DL, Appelo CAJ. 2013. Description of input and examples for PHREEQC version 3 – A computer program for speciation, batch-reaction, one-dimensional transport, and inverse geochemical calculations. U.S. Geological Survey, Washington D.C., p. 497. http://pubs.usgs.gov/tm/06/a43

**Table S1** – **Number of mapped reads per sample from amplicon sequencing of *mcrA*** **and accession numbers of the corresponding samples in the NCBI Sequence Read Archive.**

| **Lake** | **Latitude** | **Longitude** | **Reads** | **Accession** |
| --- | --- | --- | --- | --- |
| Achtriochtan | 56.667 | -5.033 | 35562 | SAMN44672983 |
| Awe | 58.095 | -4.974 | 24206 | SAMN44672979 |
| Ba | 56.608 | -4.755 | 57960 | SAMN44672984 |
| Claise | 58.370 | -5.078 | 56427 | SAMN44672976 |
| Kinardochy | 56.671 | -4.001 | 17432 | SAMN44672971 |
| Stack | 58.323 | -4.911 | 28795 | SAMN44672974 |
| Allt Ceann | 58.304 | -4.902 | 22268 | SAMN44672973 |
| Assynt | 58.151 | -4.982 | 17318 | SAMN44672978 |
| Bassenthwaite | 54.669 | -3.241 | 92371 | SAMN44672986 |
| Lochy | 56.992 | -4.864 | 47762 | SAMN44672982 |
| Lochy | 57.025 | -4.825 | 55676 | SAMN44672981 |
| Lomond | 56.147 | -4.659 | 58837 | SAMN44672985 |
| Lubnaig | 56.293 | -4.294 | 24152 | SAMN44672970 |
| Ness | 57.331 | -4.452 | 19854 | SAMN44672980 |
| Shin | 58.145 | -4.605 | 25638 | SAMN44672972 |
| Stack | 58.345 | -4.957 | 19917 | SAMN44672975 |
| Stack | 58.334 | -4.929 | 37616 | SAMN44672977 |
| Tay | 56.526 | -4.120 | 33323 | SAMN44672969 |
| Ullswater | 54.569 | -2.930 | 47947 | SAMN44672987 |

**Table S2** – **Model comparison statistics for P1 to P18 presented in the Main Text**. Models tested different predictions for how sediment chemistry, including percent organic matter (OM) in sediment, sediment polyphenolic concentrations (polyph), and microbial communities, including abundances (abund) of methanogen (*mcrA*) and methanotroph (*mmoX* or *pmoA*) marker genes in sediment, diversity (divers) of *mcrA*, and summed relative abundance of individual methanogens associated with methanogenesis (comp) influenced CH_4_ fluxes. We modified predictions *P1*-*P6* to allow for different covariation in sediment chemistry associated with separate unmeasured processes denoted by *U*_1_ to *U_4_*. For each model, we report the Akaike information criterion corrected for small sample sizes (AICc), number of estimated parameters (K) and model weight (percentage support out of candidate set). Bolded prediction is the best supported of the candidate set, i.e. lowest AICc by >2.

| **Prediction** | **CH_4_ model** | **Chemistry covariation** | **AICc** | **K** | **Weight** |
| --- | --- | --- | --- | --- | --- |
| *P1* electrochemical | %OM → polyph → redox → CH_4_ | N/A | 939.3 | 49 | 0.8 |
| ***P2* electroselective** | **%OM → polyph → redox → mcrA comp → CH_4_** | **N/A** | **929.6** | **49** | **98.5** |
| *P3* polyphenolic substrate/chelator | %OM → polyph → CH_4_ |  | 942.3 | 49 | 0.2 |
| *P4* methanogen inhibition | %OM → polyph → mcrA abund → CH_4_ | N/A | 941.9 | 49 | 0.2 |
| *P5* methanogen diversity | %OM → polyph → mcrA divers → CH_4_ |  | 941.6 | 49 | 0.2 |
| *P6* methanotroph oxidation | %OM → *mmoX* abund → CH_4_  %OM → *pmoA* abund → CH_4_ | N/A | 943.1 | 50 | 0.1 |
| *P7* electrochemical | %OM → polyph → redox → CH_4_ | %OM ← *U*_1_ → polyph  ↘ ↙  redox | 963.9 | 47 | <0.1 |
| *P8* electroselective | %OM → polyph → redox → mcrA comp → CH_4_ | %OM ← *U*_1_ → polyph  ↘ ↙  redox | 954.3 | 47 | <0.1 |
| *P9* polyphenolic substrate/chelator | %OM → polyph → CH_4_ | %OM ← *U*_1_ → polyph  ↘ ↙  redox | 967.0 | 47 | <0.1 |
| *P10* methanogen inhibition | %OM → polyph → mcrA abund → CH_4_ | %OM ← *U*_1_ → polyph  ↘ ↙  redox | 966.6 | 47 | <0.1 |
| *P11* methanogen diversity | %OM → polyph → mcrA divers → CH_4_ | %OM ← *U*_1_ → polyph  ↘ ↙  redox | 966.2 | 47 | <0.1 |
| *P12* methanotroph oxidation | %OM → *mmoX* abund → CH_4_  %OM → *pmoA* abund → CH_4_ | %OM ← *U*_1_ → polyph  ↘ ↙  redox | 967.7 | 48 | <0.1 |
| *P13* electrochemical | %OM → polyph → redox → CH_4_ | *U*_2_ *U*_3_ *U*_4_  ↙ ↘ ↙ ↘ ↙  %OM redox polyph | 983.3 | 46 | <0.1 |
| *P14* electroselective | %OM → polyph → redox → mcrA comp → CH_4_ | *U*_2_ *U*_3_ *U*_4_  ↙ ↘ ↙ ↘ ↙  %OM redox polyph | 973.6 | 46 | <0.1 |
| *P15* polyphenolic substrate/chelator | %OM → polyph → CH_4_ | *U*_2_ *U*_3_ *U*_4_  ↙ ↘ ↙ ↘ ↙  %OM redox polyph | 986.3 | 46 | <0.1 |
| *P16* methanogen inhibition | %OM → polyph → mcrA abund → CH_4_ | *U*_2_ *U*_3_ *U*_4_  ↙ ↘ ↙ ↘ ↙  %OM redox polyph | 985.9 | 46 | <0.1 |
| *P15* methanogen diversity | %OM → polyph → mcrA divers → CH_4_ | *U*_2_ *U*_3_ *U*_4_  ↙ ↘ ↙ ↘ ↙  %OM redox polyph | 985.6 | 46 | <0.1 |
| *P18* methanotroph oxidation | %OM → *mmoX* abund → CH_4_  %OM → *pmoA* abund → CH_4_ | *U*_2_ *U*_3_ *U*_4_  ↙ ↘ ↙ ↘ ↙  %OM redox polyph | 987.0 | 47 | <0.1 |

**Table S3 – Low abundance of archaeal anaerobic methanotrophs (ANMEs)**. The number of *mcrA* reads mapped to each of five Genbank accessions (columns) associated with ANME was relatively low. Of these accessions, HQ651881, HQ651880 and HQ341608 were from an unpublished study of archaea and not confirmed as ANMEs. Similarly, KF758448 and KF758447 were unidentified archaea presumed to be part of the ANME-2d clade. Only KX290023 was classified as an ANME (*Candidatus Methanoperedens* sp.).

| **Lat** | **Lon** | | **Lake** | **HQ651881** | **HQ651880** | **HQ341608** | **KF758448** | **KF758447** | **KX290023** |
| --- | --- | --- | --- | --- | --- | --- | --- | --- | --- |
|  | | *Sites with macrophyte vegetation* | | | | | | | |
| 56.667 | -5.033 | | Achtriochtan | 4598 | 0 | 0 | 0 | 0 | 0 |
| 58.095 | -4.974 | | Awe | 322 | 6 | 4 | 0 | 0 | 0 |
| 56.608 | -4.755 | | Ba | 5894 | 16 | 0 | 0 | 0 | 0 |
| 58.370 | -5.078 | | Claise | 1294 | 68 | 1 | 0 | 0 | 0 |
| 56.671 | -4.001 | | Kinardochy | 1234 | 2 | 0 | 0 | 0 | 0 |
| 58.323 | -4.911 | | Stack | 1627 | 1 | 0 | 0 | 0 | 0 |
|  | | *Sites without macrophyte vegetation* | | | | | | | |
| 58.304 | -4.902 | | Allt Ceann | 1919 | 0 | 0 | 0 | 0 | 0 |
| 58.151 | -4.982 | | Assynt | 727 | 0 | 48 | 0 | 0 | 0 |
| 54.669 | -3.241 | | Bassenthwaite | 4596 | 1 | 6 | 1 | 0 | 3 |
| 57.025 | -4.825 | | Lochy | 2844 | 6 | 0 | 0 | 0 | 0 |
| 56.992 | -4.864 | | Lochy | 2529 | 0 | 0 | 0 | 0 | 0 |
| 56.147 | -4.659 | | Lomond | 3045 | 1 | 8 | 0 | 0 | 0 |
| 56.293 | -4.294 | | Lubnaig | 3297 | 0 | 0 | 0 | 0 | 0 |
| 57.331 | -4.452 | | Ness | 2 | 0 | 0 | 11 | 8 | 1 |
| 58.145 | -4.605 | | Shin | 74 | 17 | 0 | 0 | 0 | 0 |
| 58.345 | -4.957 | | Stack | 2098 | 0 | 0 | 0 | 0 | 0 |
| 58.334 | -4.929 | | Stack | 2237 | 0 | 0 | 0 | 0 | 0 |
| 56.526 | -4.12 | | Tay | 2708 | 0 | 0 | 0 | 0 | 0 |
| 54.569 | -2.93 | | Ullswater | 6465 | 0 | 1 | 0 | 0 | 0 |

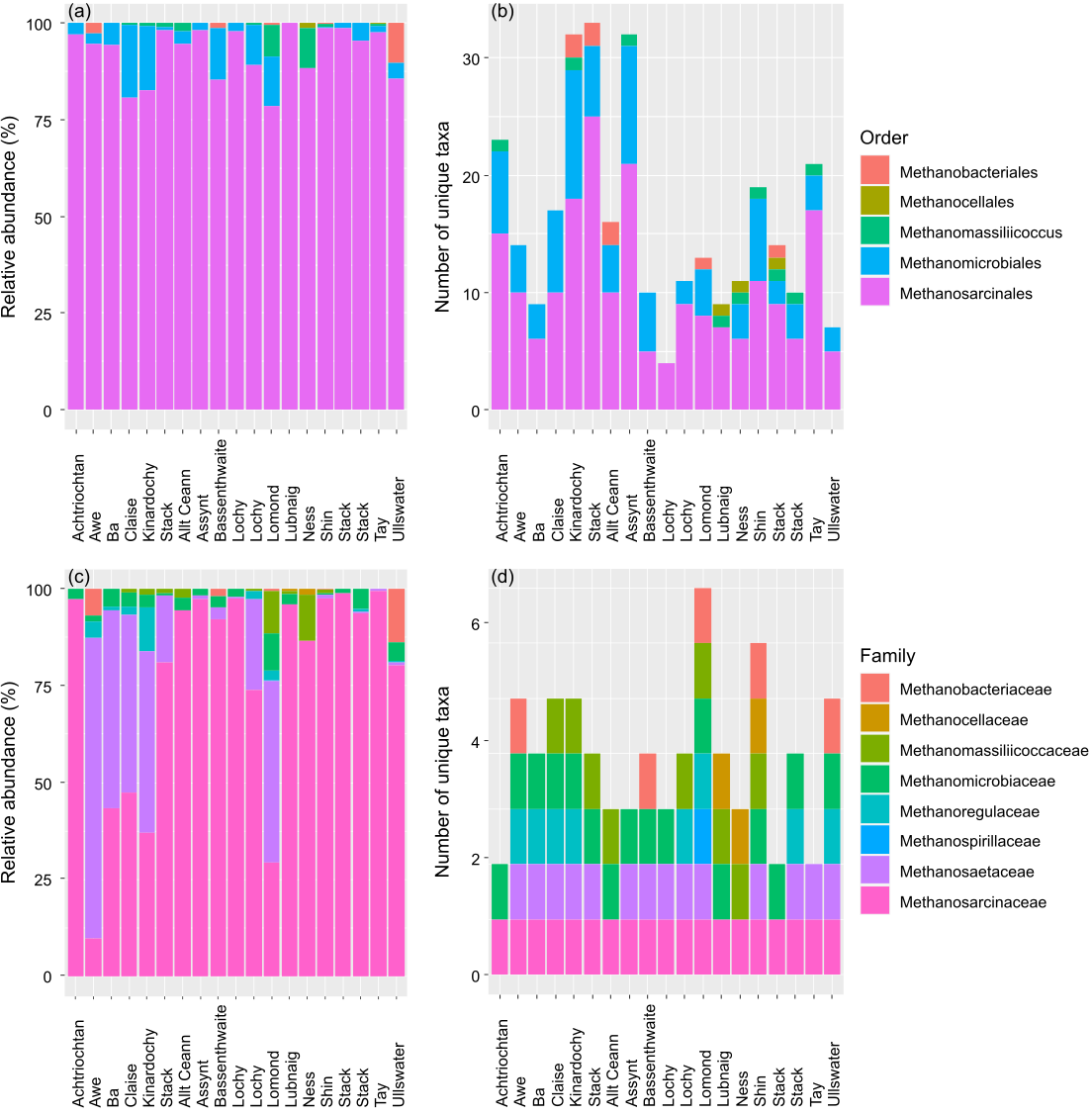

**Figure S1. Methanogen community composition is dominated by methylotrophic and acetoclastic taxa**. We calculated the (a) relative abundance and (b) number of unique taxa for *mcrA* reads that could be mapped at least to the Order-level in each of 19 sampling sites. Sampling sites are organised identically as in Table 1. Methylotrophic methanogenesis is likely facultative in Methanosarcinales and Methanobacteriales, whereas Methanomassiliicoccus are likely obligate methylotrophs (Ellenbogen and others 2023). Acetoclastic methanogenesis can be found in Methanosarcinales (Liu and Whitman 2008; Ou and others 2022).

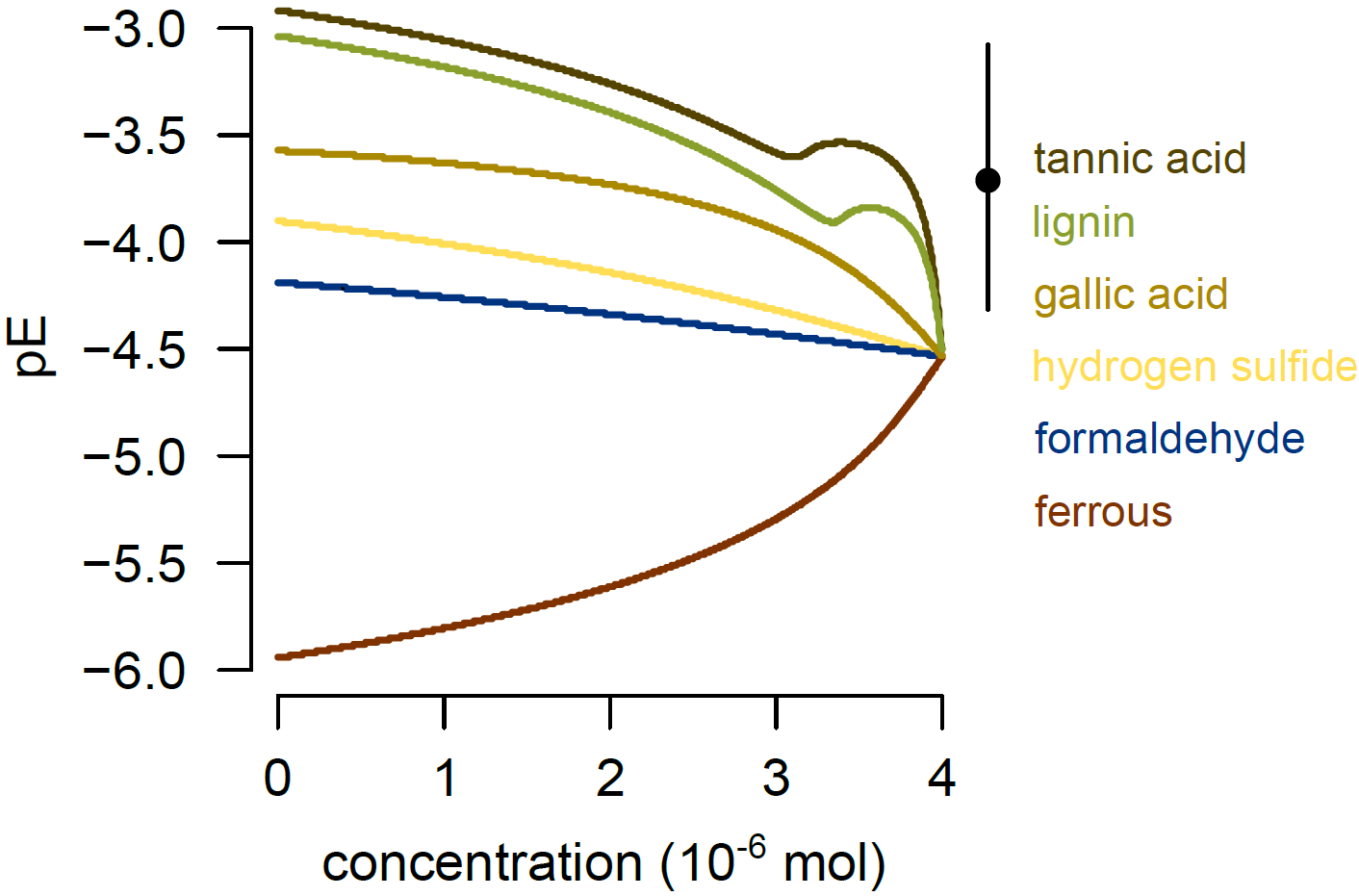

**Figure S2. Polyphenolics reduce redox more than other compounds**. We calculated redox potential pE (i.e., negative log of electron activity) when dissolving six common reducing agents into lake water using the geochemical model pH-redox-equilibrium (PHREEQC) version 3 (Parkhurst and Appelo 2013). In the absence of detailed water chemistry data for each study site, we chose a representative solute composition from Loch Morlich (latitude: 57.17, longitude: -3.7) given in Livingstone (1963). We dissolved tannic acid (C_76_H_52_O_46_), lignin (C_81_H_92_O_28_), gallic acid (C_7_H_6_O_5_), hydrogen sulfide (H_2_S), formaldehyde (CH_2_O; how PHREEQC typically represents organic matter; Parkhurst and Appelo 2013), and ferrous (Fe^2+^). Although there is no way to determine concentrations of polyphenolics degraded in our field observations (i.e., our measurements represented the outcome of decomposition), we tested how pE changed when between 0 to 100% of polyphenolics were degraded from an initial concentration of 4 × 10^-6^ mols. This initial concentration was chosen because it was representative of our field data (Table 1), and we plotted the concentration remaining after degradation on the *x*-axis to be comparable with our measurements. Plotted point is the mean ± standard error for pE observed in Table 1.
